# Compartmentalized sesquiterpenoid biosynthesis and functionalization in the *Chlamydomonas reinhardtii* plastid

**DOI:** 10.1101/2024.08.21.608953

**Authors:** Sergio Gutiérrez, Sebastian Overmans, Gordon B. Wellman, Kyle J. Lauersen

## Abstract

Terpenoids play key roles in cellular metabolism, with some organisms having evolved expanded terpenoid profiles for specialized functions such as signaling and defense. Many terpenoids have applications in pharmaceuticals, fragrances, and agriculture, but their harvest from natural sources can be challenging. Heterologous production of specialty terpenoids in microbial hosts offers an alternative, using terpene synthases and further enzymatic decoration to expand chemical complexity and functionality. Here, we explored the heterologous production of 10 different sesquiterpenoids (STPs, C_15_) and their further biofunctionalization mediated by cytochrome P450s (CYPs) in the green alga *Chlamydomonas reinhardtii*. STP synthases were expressed from the nuclear genome and localized to the algal plastid, coupled with co-expression of selected CYPs. STP production in the plastid was supported by farnesyl pyrophosphate synthase fusions to STP synthases, and CYPs were modified for soluble localization in the plastid stroma by removing transmembrane domains. Various target CYPs were screened for STP functionalization in the alga, and different product ratios were generated based on trophic modes. Overall STP yields ranged between 250-2500 µg L^−1^ under screening conditions, with CYP-mediated functionalization reaching up to 80% of accumulated heterologous STP products. Living two-phase terpenoid extractions with different perfluorinated solvents revealed variable performances based on sesquiterpenoid functionalization and solvent type. This work demonstrates the feasibility of generating heterologous functionalized terpenoid products *in alga* using soluble, plastid-localized CYPs without reductase partners. However, overall improvements in photobioreactor cultivation concepts will be required to facilitate the use of algal chassis for scaled production.

**Significance Statement:** This study demonstrates the feasibility of producing and modifying heterologous terpenoid products in the algal plastid using sesquiterpene synthases (STPS) and soluble cytochrome P450s (CYPs). We show that algal cultivation conditions influence the composition and ratios of functionalized terpenoid products, which can be extracted through solvent-based ‘milking’ during growth. The reducing equivalents that enable CYP activity in the plastid appear derived from photosynthetic electrons without requiring the co-expression of a cytochrome P450 reductase (CPR) partner, simplifying engineering strategies. As algae can be cultivated with minimal inputs (trace elements, light, and CO_2_) from sources like wastewater, this approach offers the potential for sustainable production of complex specialty terpenoid chemicals.

## 1. Introduction

Terpenoids are one of the most diverse classes of natural organic compounds, playing crucial roles in biological processes across all domains of life ^1, 2^. These molecules function in photoprotection, photosynthesis, electron transport, defense, and signaling ^1, 3, 4^. Terpenoids have wide-ranging applications in medicine, flavoring, and fragrances ^4, 5^. However, their structural complexity poses challenges for chemical synthesis ^2, 6^. Consequently, terpenoid production often relies on extraction from plant sources, resulting in low yields and impurities ^3, 6^. Harvesting terpenes from native organisms presents additional issues, as natural sources cannot meet demand due to slow growth rates or cultivation difficulties ^2, 3^, and excessive harvesting can cause ecological disruption ^4^. To address these issues, biotechnological approaches through metabolic engineering of microbes have been explored as alternatives for terpenoid production ^6^. While conventional strategies employ fermentative microorganisms, photosynthetic microalgae offer unique advantages due to their native terpenoid biosynthesis pathways and light-driven metabolism ^6–8^.

Terpenoid biosynthesis begins with the formation of five-carbon (C_5_) isoprene units: isopentenyl pyrophosphate (IPP, C_5_) and its isomer dimethylallyl pyrophosphate (DMAPP, C_5_) ^2^. These units are generated through the mevalonate (MVA) pathway or the 2-C-methyl-D-erythritol 4-phosphate (MEP) pathway ^1, 2^. Prenyltransferases catalyze the sequential addition of IPP and DMAPP to produce linear precursors, including geranyl pyrophosphate (GPP, C_10_), farnesyl pyrophosphate (FPP, C_15_), and geranylgeranyl pyrophosphate (GGPP, C_20_) ^2^. Terpenoid synthases (TPS) convert these precursors into cyclic terpenoid skeletons through complex reactions ^6^. This process yields various terpenoid (C_20_), triterpenoids (C_30_), and tetraterpenoids (C_40_) ^9^. The resulting skeletons often undergo further modification through functionalization catalyzed by enzymes like acetyltransferases, carboxylases, and cytochrome P450 (CYP) monooxygenases ^2, 8^. CYPs are found overexpressed in tissues where terpenoids accumulate, allowing efficient substrate access and interaction with cofactors and redox partners ^10^.

This study investigates heterologous complex terpenoid biosynthesis in *Chlamydomonas reinhardtii* by localizing sesquiterpene production and functionalization reactions in the algal plastid. We leverage advances in transgene design for robust expression, native terpenoid precursor supply in the photosynthetic cell, and the redox environment of its plastid to generate heterologous terpenoids and chemically functionalize them. By integrating terpenoid biosynthesis and CYP-mediated functionalization into the plastid, we aimed to leverage its chemical reduction potential to mediate functional group addition to heterologous STPs without the expression of partnering cytochrome P450 reductases (CPRs). We focused on fragrant sesquiterpenoids (STP, C_15_) derived from agarwood and sandalwood, where minimal chemical modifications to the terpenoid backbone can expand scent profiles. We evaluated the effects of carbon sources on product formation and examined the behavior of produced terpenoids when algal cultures interact with different extractants. By exploring *C. reinhardtii* as a host for compartmentalized terpenoid production, this work contributes to the development of light-driven methods for producing valuable chemical compounds.

## 2. Results and Discussion

### 2.1. Sesquiterpenoid production from terpene synthases located in the cytoplasm or chloroplast

*C. reinhardtii* has shown promise for photosynthetic production of non-native terpenoids ^7, 11–16^. This microalga produces all terpenoid precursors for photosynthesis and other cellular processes using only the MEP pathway in its plastid ^6^. Recent advances in transgene design have improved the expression of nuclear-encoded transgenes, facilitating metabolic engineering efforts for non-native terpene production and elucidating the metabolic flexibility of the photosynthetic cell ^16, 17^. Although *C. reinhardtii* lacks endogenous sesquiterpenoid (STP) synthases, it can be engineered to produce diverse STPs by redirecting carbon flux from cytosolic farnesyl pyrophosphate (FPP) ^12, 14, 15, 17–20^. In contrast, FPP levels in the plastid are natively low but can be increased to produce heterologous STPs by overexpressing a native or non-native FPP synthase localized to this subcellular compartment ^12, 15, 18, 20^.

We compared the yields of 10 sesquiterpene synthases (STPSs) expressed from the algal nuclear genome and localized to either the cytoplasm or the plastid. Plastid-targeted constructs were designed with C-terminal FPPS fusions, as previous studies have shown that free FPP levels in the plastid are minimal and FPPS fusion to STPSs does not enhance productivity in the cytoplasm ^14, 20^. The cell line employed features a constitutive knockdown of the native squalene synthase, which competes directly for cytoplasmic FPP with the introduced STPSs^12^. This modification allowed us to compare yield variations under best-case production scenarios between the cytoplasmic STPSs and plastid-localized STPSs fused to FPPS. We evaluated three different construct designs (**Fig. 1A–C, SI Appendix, Table S1, File S1, and Fig. S1–S3**): a cytoplasmic targeted STPS alone and two different plastid-targeted FPPS fusions to STPSs, either the from *Saccharomyces cerevisiae* (Erg20) or *Escherichia coli* (ispA). Many sesquiterpenoids were successfully produced, including aristolochene, valencene, selinene, santalene, bisabolol, τ-cadinol, α-cadinene, β-cadinene, γ- cadinene, murolene, α-guaiene, β-guaiene, δ-guaiene, α-humulene, alloaromadendrene, valerianol, and patchoulol (**Fig. 1D, SI Appendix, Files S2– S4**) in both compartments, with yields ranging from ∼250 to 2500 µg L^−1^ (**Fig. 1E, SI Appendix, Table S2, and Fig. S5**). The yields of STPSs were comparable for each sesquiterpene product from both cytoplasm (STPS alone) and plastid-targeted (STPS-FPPS fusion) containing transformants (**Fig. 1E, SI Appendix, Table S2, and Fig. S5**). Plastid-localized bisabolol and cadinol synthases achieved higher levels of bisabolol and cadinol production than their cytoplasmic counterparts. Bisabolol (construct C07) reached 84 µg L^−1^, while cadinol (construct C08) produced 208 µg L^−1^ in (**Fig. 1E, SI Appendix, Table S2, and Fig. S5).**

**Fig. 1.**
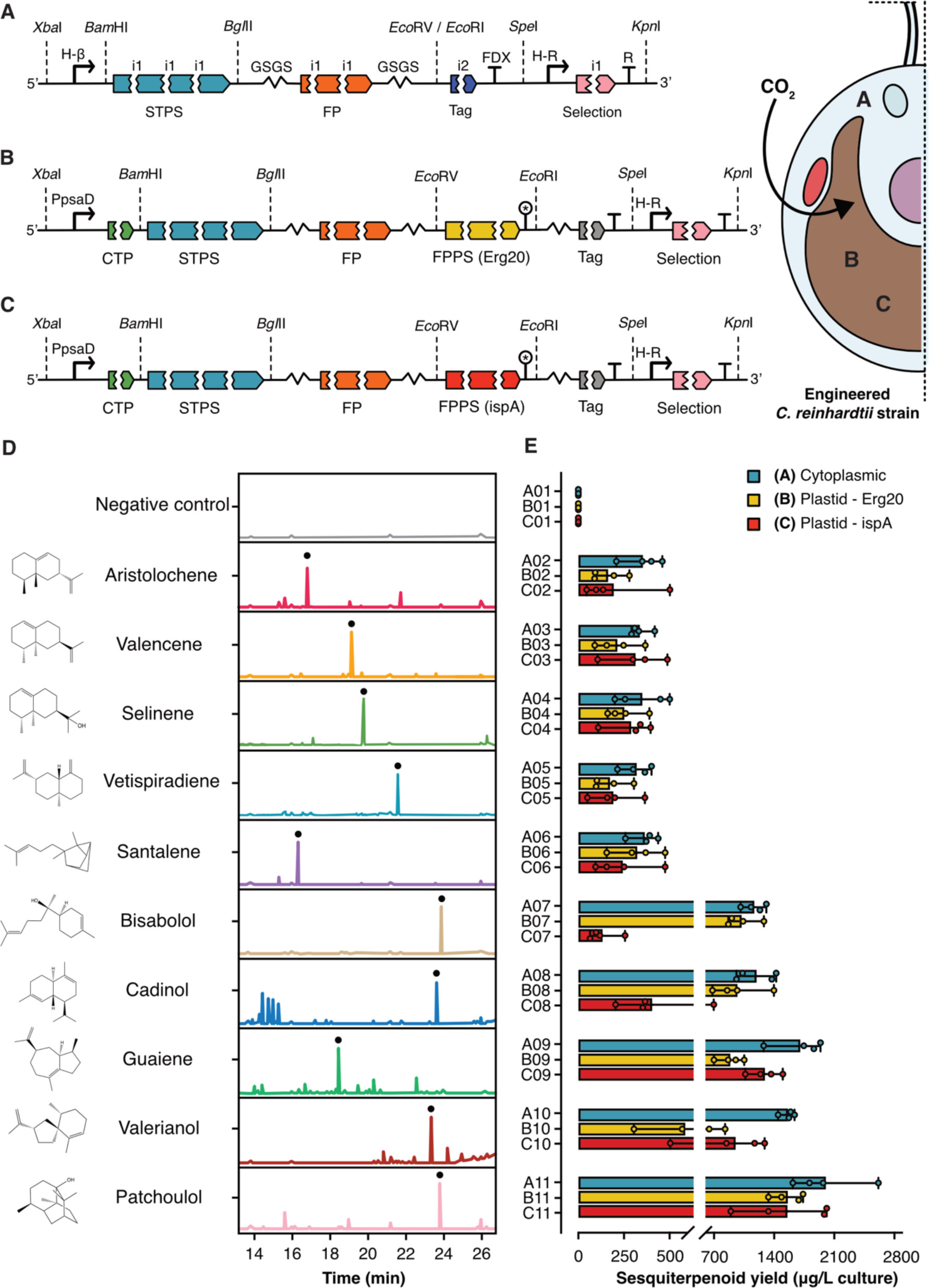
Genetic constructs and sesquiterpenoid production in engineered *C. reinhardtii* strains. (A-C) Schematic representation of genetic constructs for sesquiterpenoid production in *C. reinhardtii*. Constructs include sesquiterpene synthases (STPS) fused to fluorescent protein reporters (FP) and farnesyl pyrophosphate synthases (FPPS) targeted to: (**A**) cytoplasm, (**B**) plastid with *Erg20* FPPS (*S. cerevisiae*), and (**C**) plastid with *ispA* FPPS (E. coli). Promoters: H-ß (heat-shock protein/beta-tubulin), pPsaD (photosystem I subunit II promoter), H (heat-shock protein 70S promoter), R (RuBisCO small subunit 2 promoter). CTP: chloroplast transit peptide (PsaD). RBCS intron 1 (i1) and intron 2 (i2) are spread throughout the coding sequences of optimized genes. FDX: ferredoxin 1 terminator. Inset: Engineered *C. reinhardtii* strain with chloroplast (brown), indicating modified carotenoid synthesis; letters indicate intended localization of recombinant enzyme products. (**D**) Chromatograms of sesquiterpenoid products from each STPS expression, compared to a parental strain negative control extract. Black dots indicate intended sesquiterpenoid products. (**E**) Sesquiterpenoid yields (µg/L culture) for each construct. Data for genetic constructs and GC-MS/FID can be found in **SI Appendix Tables S1–S3 and Files S1–S4**.

The choice of FPPS influenced production efficiency in an unpredictable manner for each STPS. We found that ispA fusion constructs **C02-C11** (**Fig. 1E; SI Appendix, Table S2, Fig. S3, and S5**) yielded higher titers for aristolochene, valencene, selinene, guaiene, and valerianol than their Erg20 counterparts (**B02-B11; Fig. 1E; SI Appendix, Table S2, Fig. S2, and S5**). These results suggest that selecting the appropriate FPPS can enhance the production of specific STPs; however, iterative empirical testing for each target STPS-FPPS fusion is required ^21, 22^. Next, we investigated whether the reducing environment of the plastid could mediate the chemical functionalization of heterologous STP products by co-expressing CYPs specific to these compounds.

### 2.2. Functionalization of heterologous sesquiterpenoids mediated by co-expression of plastid-targeted P450s

In native hosts, terpenoids undergo chemical modifications that enhance their complexity and biological activities ^14, 23^. Metabolic engineering can recapitulate these reactions by expressing the corresponding metabolic pathway in a foreign host ^2, 4, 9, 24, 25^. Cytochrome P450 monooxygenases (CYPs) commonly catalyze reactions that add hydroxyl or other functional groups to terpenoids. CYPs receive electrons from CPRs, typically on the cytoplasmic side of the endoplasmic reticulum (ER) membrane ^10, 19, 26^. CYPs and CPRs contain transmembrane (TM) anchors and have been successfully expressed and shown to function in non-native hosts such as yeasts and tobacco ^10, 11, 27^. Recent studies have demonstrated that when CYPs are expressed and localized in cyanobacteria or plant plastids, photosynthesis-derived electrons can replace CPRs in driving CYP reactions ^27–31^.

As we produced reasonable amounts of STPs in the *C. reinhardtii* plastid by introducing STPS-FPPS fusions (**Fig. 1E, SI Appendix, Table S2, and Fig. S2– S3**), we aimed to use the redox potential of the plastid for their chemical functionalization ^10, 11, 19^. For each STP studied, we co-expressed and targeted CYPs to the algal plastid that had predicted or were previously shown to be responsible for mediating their functionalization (**SI Appendix, Table S1, File S1, Fig. S2, and S4**). Transformants confirmed to express both STPS-FPPS fusion and heterologous CYPS were cultivated with solvent overlays, and the products were analyzed by GC-MS/FID. Changes in chromatogram peaks were observed for all strains with co-expressed CYPs compared to those expressing STPSs alone (**Fig. 2A, SI Appendix, Table S3, and Files S5–S6**). These changes indicate the presence of sesquiterpenoid derivatives, confirming the successful modification of base STPs in the plastid through CYP co-expression. Functionalization efficacy varied among CYPs, with some producing the expected product for a specific STP, while others generated numerous side products or inefficiently synthesized target compounds (**Fig. 2A, 2B, SI Appendix, Tables S7–S8, and Files S5–S6**). As no CPR was co-expressed in these strains, the results indicate that electrons present in the plastid can drive these targeted chemical modifications of heterologous terpenoids ^10, 11, 14, 19, 25, 27^. The exact electron donor is unknown, but it could be ferredoxin or simply the reducing environment of the plastid itself in illuminated conditions.

**Fig. 2.**
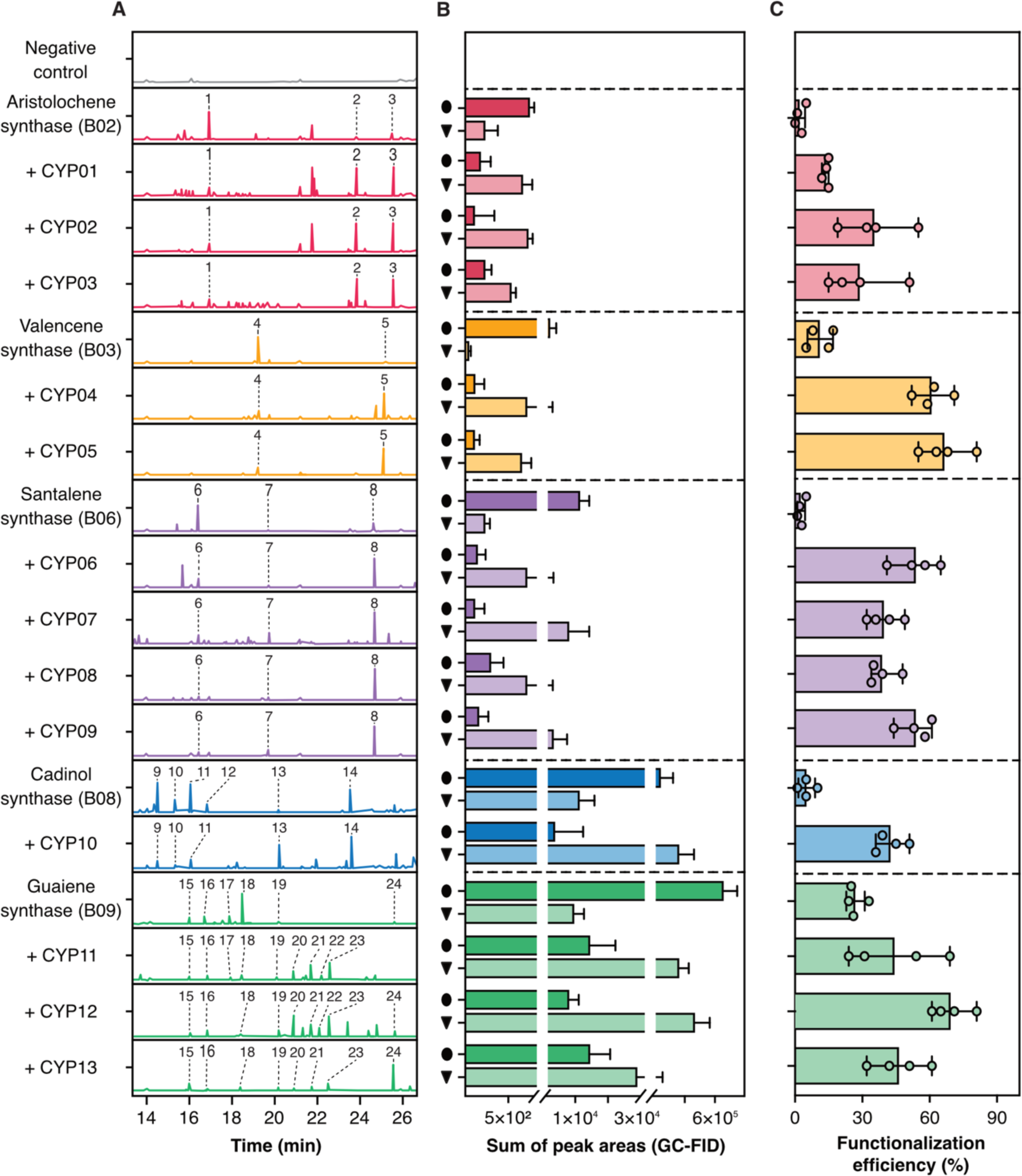
Plastid-targeted sesquiterpenoid biosynthesis and functionalization by cytochrome P450 enzymes in *C. reinhardtii*. Sesquiterpene synthases (STPS) and cytochrome P450 enzymes (CYPs) were targeted to the algal chloroplast. Each STPS construct included C-terminal ScErg20 FPPS, while CYPs were co-expressed through plastid targeting and without transmembrane domains. **(A)** GC-MS chromatograms of dodecane extracts from *C. reinhardtii* strains expressing plastid localized STPS and CYP combinations, numbers indicate specific compounds identified by MS. **(B)** Quantitative analysis of sesquiterpenoid production based on GC-FID data. Circles: sum of peak areas for sesquiterpenoids; triangles: sum of peak areas for modified sesquiterpenoids. Data represent mean ± SD (n=12, 4 transformants × 3 biological replicates). **(C)** Functionalization efficiency (%) of each CYP, calculated as the fraction of functionalized sesquiterpenoids from total sesquiterpenoids. Circles represent individual transformants. Compounds: [1] Aristolene, [2] Aristolochene, [3] Aristolochone, [4] Valencene, [5] Nootkatone, [6] α-Santalene, [7] Bergamotol, [8] Santalol, [9] α-Cadinene, [10] δ-Cadinene, [11] β-Cadinene, [12] Muurolene, [13] Muurolol, [14] τ-Cadinol, [15] α-Guaiene, [16] β-Guaiene, [17] α-Humulene, [18] δ-Guaiene, [19] Alloaromadendrene, [20] α-Guaiol, [21] β-Guaiol, [22] Globulol, [23] Rotundone, [24] Alloaromadendrene oxide. GC-MS/FID data in **SI Appendix File S5, Tables S3, and Fig. S6–S7.**

For each STPS-CYP combination, we identified one CYP which generated the target functionalization as predicted or desired (**Fig. 2B, 2C, SI Appendix, Tables S4, and S7–S8**). Functionalization efficiency, calculated as the total sum of new peaks relative to the original STPS abundance, varied among each (**Fig. 2C, SI Appendix, Fig. S7, Tables S3, and S5**). We observed the following conversions: (1) Aristolochene to aristolochone by CYP02 (UniProt: W6QP06) at 45 ± 21% efficiency (mean ± SD). (2) Valencene to nootkatone by CYP04 (UniProt: E1B2Z9) at 58 ± 18% efficiency. (3) Santalene to bergamotol and santalol by CYP09 (UniProt: VR5EU4) at 55 ± 13% efficiency. (4) α-Cadinene, β-cadinene, γ-cadinene, and muurolene to muurolol and α-cadinol by CYP10 (UniProt: A0A0N9H930) at 51 ± 8% efficiency. (5) α-guaiene, β-guaiene, δ-guaiene, α-humulene, and alloaromadendrene to α-guaiol, β-guaiol, globulol, rotundone, and alloaromadendrene oxide by CYP12 (UniProt: E3W9C4) at 66 ± 19% efficiency.

Selecting appropriate CYPs to ensure product formation remains challenging, and our screening suggests that empirical testing is necessary to determine the correct combination (**Fig. 3**)^10, 12, 19, 27^. Each CYP also led to the formation of off-target products **(Fig. 3A, SI Appendix, Tables S3–S4, and S7–S8**). A heatmap of the major identifiable products illustrates the chemical diversity obtained with each CYP on tested STPs (**Fig. 4, SI Appendix, Fig. S6, and Table S7–S8**). Functionalization efficiency consistently remained below 80% (**Fig 2C, SI Appendix, Table S4, and Fig. S7**), indicating room for improvement, which may be attained in the future through the formation of metabolons or artificial STPS-CYP associations via enzyme engineering strategies ^10, 11, 24, 25, 28–31^.

**Fig. 3.**
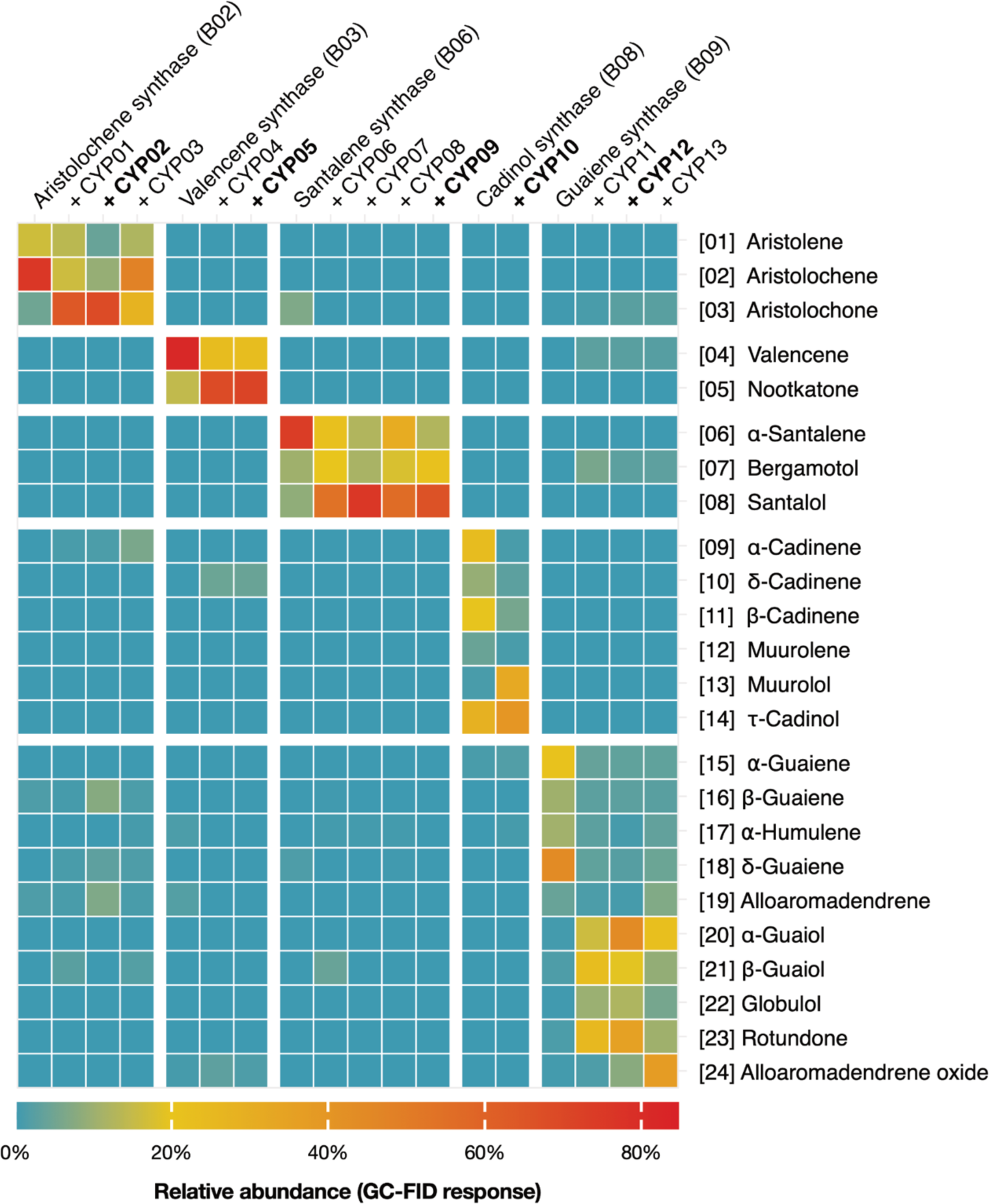
Relative abundance of sesquiterpenoids produced in *C. reinhardtii* with different cytochrome P450s (CYPs). Heat map showing the relative abundance of sesquiterpenoid compounds based on GC-FID response. Columns represent sesquiterpene synthase and CYP combinations. Color intensity indicates relative abundance: red (high) to blue (low). Sesquiterpenoid compounds identified by MS are listed on the right with corresponding numbers. GC-MS/FID data in **SI Appendix File S5, Table S3, and Fig. S6**.

**Fig. 4.**
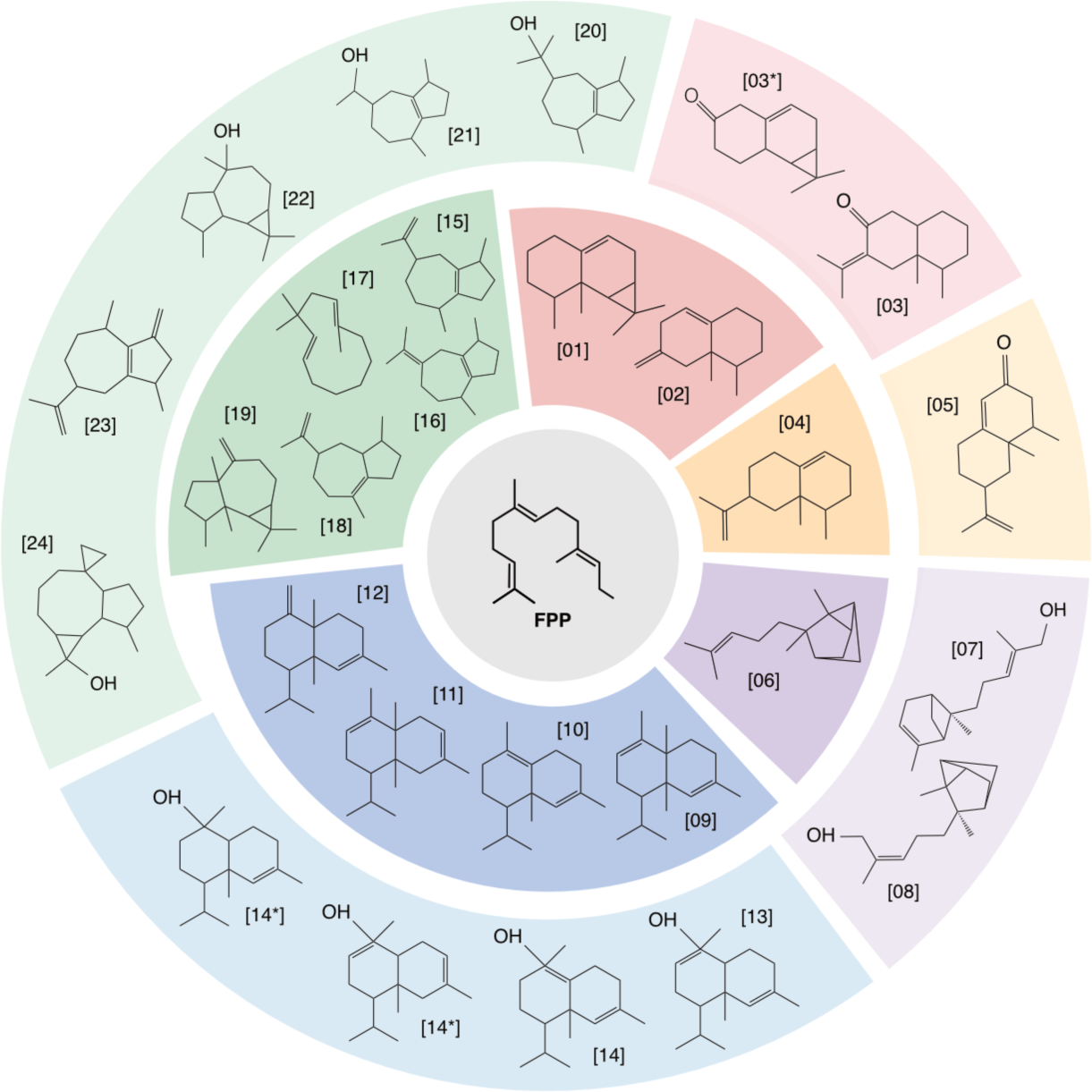
Sesquiterpenoid compounds produced from farnesyl pyrophosphate (FPP) from the *C. reinhardtii* plastid. Illustration of sesquiterpenoid chemical structures identified in this work. Colored sections depict distinct sesquiterpenoid classes produced through the action of specific sesquiterpene synthases and cytochrome P450 enzymes. Compounds are labeled with unique identifiers corresponding to their structures: Aristolene [01], Aristolochene [02], Aristolochone [03], Valencene [04], Nootkatone [05], α-Santalene [06], Bergamotol [07], Santalol [08], α-Cadinene [09], δ-Cadinene [10], β-Cadinene [11], Muurolene [12], Muurolol [13], τ-Cadinol [14], α-Guaiene [15], β-Guaiene [16], α-Humulene [17], δ-Guaiene [18], Alloaromadendrene [19], α-Guaiol [20], β-Guaiol [21], Globulol [22], Rotundone [23], and Alloaromadendrene oxide [24]. GC-MS/FID data in **SI Appendix File S5, Tables S7–S8**.

### 2.3. Carbon source effects on plastid sesquiterpenoid biosynthesis and functionalization

*Chlamydomonas* can grow on organic acetic acid, inorganic CO_2_, or both as carbon sources. The trophic mode of cultivation induces major rearrangements in cell architecture, prompting us to investigate the effects of these changes on the product profiles of our engineered strains ^32^. Using the most effective STPS-CYP pairs, we analyzed products from solvent milking of strains grown under three illuminated conditions: CO_2_ alone, acetate alone, or combined CO_2_+acetate (**Fig. 5**). Chromatograms revealed distinct STP and derivative profiles under these cultivation modes (**Fig. 5, SI Appendix, File S6**). Relative abundance data showed variations in sesquiterpenoid production and functionalization efficiency depending on the carbon source (**Fig. 5, SI Appendix, File S6**).

**Fig 5.**
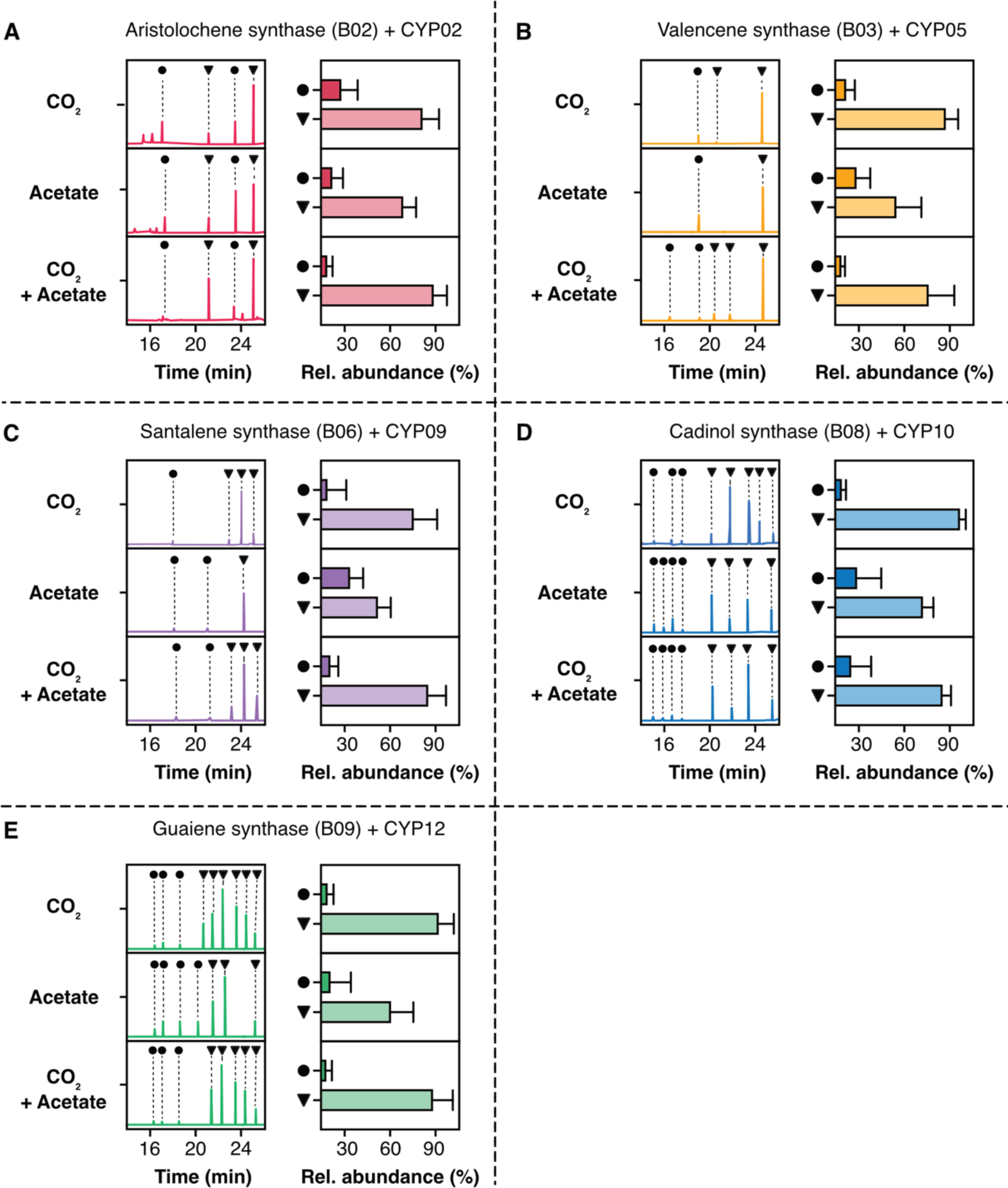
Carbon source effects on plastid-targeted sesquiterpenoid biosynthesis and functionalization in *C. reinhardtii.* GC-MS/FID analysis of dodecane overlay samples for sesquiterpenoid species accumulated by *C. reinhardtii* expressing different STPSs (B02, B03, B06, B08, B09) and corresponding cytochrome P450s (CYP02, CYP05, CYP09, CYP10, CYP12) when cultivated with three carbon source conditions: CO₂, acetate, or CO₂+acetate. Black dots represent sesquiterpenoid compounds; black triangles indicate functionalized derivatives. Relative abundance plots quantify production levels under each condition. **(A)** Aristolochene synthase (B02) + CYP02; **(B)** Valencene synthase (B03) + CYP05; **(C)** Santalene synthase (B06) + CYP09; **(D)** Cadinol synthase (B08) + CYP10; **(E)** Guaiene synthase (B09) + CYP12. For each panel, chromatograms (left) show product retention times; bar graphs (right) depict relative abundance. GC-MS/FID data in **SI Appendix File S6, Tables S6–S8.**

Aristolochene synthase + CYP02 (**Fig. 5A, SI Appendix, File S6**) exhibited three major peaks across all conditions. CO_2_ alone and CO_2_+acetate conditions yielded higher relative abundances of the functionalized product than acetate alone. Valencene synthase + CYP05 (**Fig. 5B, SI Appendix, File S6**) primarily produced nootkatone, with two minor peaks under CO_2_ and CO2+acetate conditions. Nootkatone abundance was highest with CO_2_ alone, followed by CO_2_+acetate, and lowest with acetate alone.

Santalene synthase + CYP09 (**Fig. 5C, SI Appendix, File S6**) produced multiple peaks, representing α/β-santol, bergamotol, and precursors. CO_2_ alone yielded the highest relative abundance of functionalized products, while acetate alone showed the lowest. Cadinol synthase + CYP10 (**Fig. 5D, SI Appendix, File S6**) showed multiple peaks across all conditions. Acetate and CO_2_+acetate conditions produced higher relative abundances of functionalized products compared to CO_2_ alone. The most diverse compound array was generated by guaiene synthase + CYP12 (**Fig. 5E, SI Appendix, File S6**). CO_2_ and CO_2_+acetate conditions displayed greater peak diversity than acetate alone, with CO_2_+acetate showing the highest relative abundance of functionalized products.

These results indicate that a mixed carbon source strategy enhances sesquiterpenoid production and functionalization in *C. reinhardtii*. The combination of CO_2_+acetate generally resulted in higher relative abundances of functionalized products in growth conditions tested, likely due to higher cell densities (**Fig 5, SI Appendix, File S6)**^8, 11, 25–27, 31^. The variable peak intensities and product species observed here add complexity to the prediction of functionalized STP product outputs from engineered algal cultivation. Whether product profiles can be consistently tailored during scaled cultivations is still unknown. *Chlamydomonas* is not routinely cultivated phototrophically at scale, and research is ongoing to test methods of scaled extraction of engineered terpenoids through solvent milking ^33, 34^. Future studies should address these limitations through production scale-up and long-term cultivation experiments with variable light regimes to assess production feasibility ^6, 14, 35^.

### 2.4. Culture-solvent extraction efficiencies

Extracting non-native sesquiterpenoids from *C. reinhardtii* requires the culture to grow in contact with a biocompatible solvent ^5, 21, 23, 26, 31, 33, 34, 36–38^. The choice of extraction solvent has environmental and economic implications for bioprocess designs ^39^. While dodecane is a standard biocompatible solvent for lab-scale terpenoid extraction and quantification, perfluorinated solvents offer advantages in safety, stability, and reusability ^12, 33, 34, 37, 38, 40, 41^. We recently reported a method to use perfluorinated solvents to extract algal-produced terpenoids ^12^. This method allowed the subsequent transfer of terpenes from perfluorinated solvent to ethanol through liquid-liquid separation, enabling recycling of the clean perfluorinated solvent to algal culture and direct use of the ethanol-terpene mixture for fragrance applications.

Here, we evaluated a larger pool of perfluorinated solvents for their capacity to accumulate heterologous and chemically complex sesquiterpenoids produced by *C. reinhardtii* compared to dodecane (**Fig. 6, SI Appendix, Table S5**). Terpenoid accumulation was lower in all fluorinated solvents than in dodecane, and extraction efficiencies varied across solvents and sesquiterpenoid compounds (**Fig. 6, SI Appendix, Table S5**). This variability is likely due to individual differences in solvent and sesquiterpenoid properties, as each solvent is unique, and the sesquiterpenoids have structural differences ^12, 21, 33, 34^. FC-40 and FC-770 demonstrated higher extraction capacities for bisabolol (32% and 31% compared to dodecane, respectively), while CFL7160 extracted aristolochene (15%) and selinene (16%) more effectively (**Fig. 6, SI Appendix, Table S5**) ^12–14, 18, 33, 34^. We observed enhanced accumulation of -OH group– containing STPs in all FCs tested (**Fig. 6, SI Appendix, Table S5**), with bisabolol, candinol, valerianol, and patchoulol accumulating at higher levels than aristolochene, valencene, selinene, vetispiradiene, and santalene.

**Fig 6.**
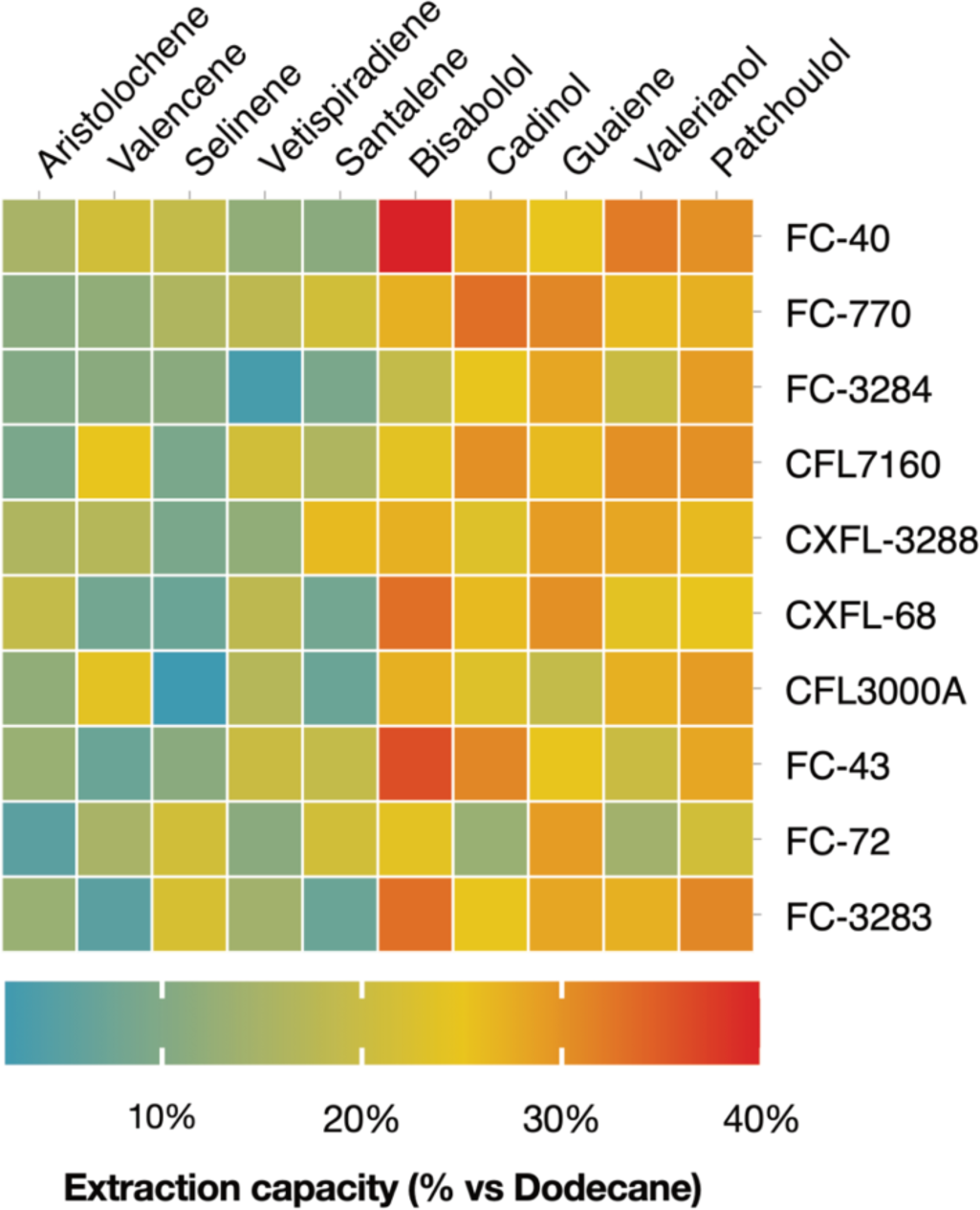
Sesquiterpenoid milking efficiencies using different perfluorinated solvents on engineered *C. reinhardtii*. Heat map comparing extraction capacities of fluorinated solvents relative to dodecane for various sesquiterpenoid compounds. Solvents tested: FC-40, FC-770, FC-3284 (perfluoro-n-dibutylmethylamine), CFL7160 (perfluoro noeny trifluoroethyl ether), CXFL-3288 (perfluorotripropylamine), CXFL-68 (perfluorotributylamine), CFL3000A (hexafluoropropene trimer), FC-43, FC-72 (perfluoro-n-dibutylmethylamine), and FC-3283 (perfluorotripropylamine). Color intensity indicates extraction capacity (blue: 0%, red: 40%). GC-MS/FID data in **SI Appendix Table S5.**

These results indicate that solvent choice will need to be tailored to individual products if scaled processes for algal-produced terpenes are to be implemented. Our screening was conducted in small volume well-plates, in which the contact surface area of perfluorinated solvents is reduced by forming liquid beads under the aqueous phase. In scaled cultivation, the surface area for solvent milking can be improved, although chemical partitioning variability is likely to persist. Scaling these extraction methods for industrial applications presents challenges for which photobioreactors have not yet been optimized ^12, 33, 36, 38^. To utilize engineered algae for heterologous terpene production, novel reactor designs that enable scalable culture-solvent interaction while maintaining optimal light-driven algae growth will be necessary.

### 2.5. Conclusions

This study demonstrates the feasibility of producing a diversity of chemically complex functionalized terpene products from the engineered algal cell. We show that functionalization of heterologous terpenoids can be achieved using soluble CYPs targeted to the plastid without the need for partner CPRs. While our investigation focused on STPs, this strategy could be applied to higher-value diterpene products as precursors for pharmaceutical production. The alga system offers a potential platform for rapid investigation of CYP activity, given the affordability of gene synthesis, quick generation of transformants within weeks, and straightforward analysis of terpenoid products. However, challenges and uncertainties remain in whether this can be scaled as a valuable production chassis for complex terpenoid compounds.

## 3. Materials and Methods

### 3.1. Algae cultivation, plasmid design, transformation, and screening

Experiments used a *C. reinhardtii* strain derived from UPN22, modified for enhanced terpenoid biosynthesis through squalene synthase knockdown and β-carotene ketolase overexpression ^5, 6, 36, 38–41^. Cultures were maintained in TAPhi-NO_3_ medium under LED illumination (150 µmol m^−2^ s^−1^). We selected ten sesquiterpene synthases (STPSs), including eight previously designed and two additional synthases - selinene synthase (UniProt: O64404) and vetispiradiene synthase (UniProt: A0A411G8M5)^12, 15^. Genes were codon-optimized and subjected to intron spreading to enable expression from the nuclear genome ^12^.

We used three pOpt3-based STPS-containing construct designs: one for cytoplasmic expression with paromomycin selection (APHVIII) and two for chloroplast-targeted expression with hygromycin selection (APHVII)^42–44^. Chloroplast-targeted STPS constructs employed the *PsaD* promoter and chloroplast targeting peptide (CTP) with mKOk fluorescent protein and either *S. cerevisiae* (Erg20) or *E. coli* (ispA) farnesyl diphosphate synthase (FPPS). As previously reported, these FPPSs included a C-terminal stop codon to maintain activity ^45^. Selected CYPs were optimized for nuclear genome expression and targeted to the algal plastid. We removed the TM domain from all CYP coding sequences to enable expression as soluble proteins in the plastid stroma. TM domains were identified through the TMHMM - 2.0 server and comparing AlphaFold models of each protein sequence to see low-structured N-terminal regions ^14, 18, 19^. CYP-*C. reinhardtii* optimized sequences were subcloned into pOpt3-based expression constructs with the *PsaD* promoter and CTP, containing the teal fluorescent protein (mTFP1) as a reporter and selection for zeocin resistance (shBle). All constructs were synthesized *de novo* and subcloned by Genscript (Piscataway, NJ, USA) (**SI Appendix, Table S1, and File S1**).

*C. reinhardtii* nuclear genome transformation used linearized plasmid DNA (restriction enzymes: *Xba*I and *Kpn*I) via a glass-bead protocol, with 10 µg DNA per transformation ^46^. After an 8-hour recovery in liquid TAPhi-NO_3_ medium ^47^ under low light, cells were plated on a medium containing spectinomycin (200 µg mL^−1^) plus paromomycin (10 µg mL^−1^), hygromycin B (15 µg mL^−1^), or zeocin (15 µg mL^−1^), individually or in combinations, matching the desired selection. Plates were exposed to continuous light for 7 days before colony selection. A PIXL colony-picking robot (Singer Instruments, UK) transferred up to 384 colonies per transformation event onto TAPhi-NO3 agar plates. After 3 days, a ROTOR robot (Singer Instruments, UK) duplicated colonies onto new plates containing amido black (150 µg mL^−1^) for fluorescence screening ^15^. We selected transformants exhibiting intense fluorescent-protein signals (**SI Appendix, Fig. S1–S4)** and transferred them to 12-well microtiter plates containing 2 mL liquid TAPhi-NO_3_ medium, cultured with agitation at 160 rpm for subsequent two-phase cultivation and solvent analysis ^12, 45^.

### 3.2. Capture of algal-produced sesquiterpenoids and their analysis by GC-FID/MS

We quantified sesquiterpenoid production using a two-phase cultivation system as described previously ^12, 33^. Four transformants were selected based on fluorescence and analyzed in triplicate. Cultures were grown in 6-well microtiter plates containing 4.5 mL TAPhi-NO_3_ medium and 500 µL dodecane overlay (10% of total volume) for 7 days ^12, 15^. To evaluate alternative perfluorocarbon (FCs) as terpenoid extraction solvents ^12, 15, 45^, we tested ten FCs alongside dodecane: CFL7160, CXFL-68, CFL3000A, CXFL-3288, FC-770, FC-3284, FC-43, FC-72, FC-40, and FC-3283 (Sigma-Aldrich, Germany; Acros Organics, Belgium; Hunan Chemfish Pharmaceutical Co., Ltd, China) (**SI Appendix, Table S5**). For these extractions, we used 1000 μL of solvent (20% of total volume) with 4 mL cultures. Dodecane formed an upper ‘overlay’ while FCs formed ‘underlays’. We quantified final culture volumes after cultivation to account for evaporation. After cultivation, the phases were separated by centrifugation at 3500 × *g* for 5 min. Both solvent fractions (FCs and dodecane) were transferred to GC vials for analysis. Cell density was measured by flow cytometry (**SI Appendix, File S7**) ^33^.

We performed GC-MS/FID analysis as described ^45^ and processed chromatograms using MassHunter software (Agilent, Germany, version B.08.00). Compounds were identified by comparing mass spectra to the NIST Mass Spectral Library (National Institute of Standards and Technology, USA). For quantification, we used calibration curves (1 – 500 μM) of purified standards in dodecane or FCs: δ-guaiene, patchoulol, α-santalene, valerianol, α-bisabolol, valencene, and cedrene (Toronto Research Chemicals, Canada) (**SI Appendix, Fig. S5**). A standard terpene mixture (MetaSci, Canada) containing 98 terpenes at 1 mM in methanol ensured accurate identification and internal library calibration (**Fig. 6, SI Appendix, Table S6**).

### 3.3. Data analysis

Each experimental condition included three biological replicates per transformant, with experiments independently repeated to ensure reproducibility. Controls comprised the parental non-transformed *C. reinhardtii* strain and vector-only constructs with respective fluorescent reporters. We performed GC-MS/FID measurements in triplicate, manually reviewing chromatograms for quality control. Terpenoid extracts were analyzed using established methods ^12, 33, 34^. We based compound identification on retention index, match factor, and comparison to the NIST library. We calculated mean production values and standard deviations for quantitative analysis and performed descriptive statistics (**SI Appendix, Tables S2–S5)**. Statistical analyses used JMP v.16 (SAS Institute, NC) and R v.3.6.2 (R Foundation for Statistical Computing, Austria). We visualized data using JMP v.16 and GraphPad Prism v.10.3 (GraphPad Software, USA). Diagrams and illustrations were created using Affinity Designer v.2.5.3 (Serif Ltd., UK), chemical structures were drawn with ChemDraw v.20.1 (PerkinElmer, MA, USA), and visual elements were integrated using Affinity Publisher v.2.5.3 (Serif Ltd., WB, UK).

## 4. Data availability

All data supporting the findings of this study are included in the article and **SI Appendix**. Source data and genetic files are available in DRYAD (https://doi.org/10.5061/dryad.zgmsbccmz). For review purposes, source data can be accessed via a temporary link (https://rb.gy/lpdr3a).

## 5. Acknowledgments

This work received support from King Abdullah University of Science and Technology competitive research grant 4715 and baseline research funding to KJL. SG and KJL would like to express their gratitude to Dr. Ahmed Alfahad of King Abdulaziz City for Science and Technology (KACST), who found and suggested several terpene synthases and cytochrome P450 enzymes used in this work.

