## Supplemental Files for "Compartmentalized sesquiterpenoid biosynthesis and functionalization in the *Chlamydomonas reinhardtii* plastid"

This PDF file includes:

Data availability
Supplementary materials and methods
Figure S1 to S5
Tables S1 to S10
Legends for Files S1 to S12
SI References

Other supporting materials for this manuscript include the following:

Files S1 to S12
Dataset (DRYAD)

### SI Appendix Content

#### Data availability

All data supporting the findings of this study are included within the article and **SI Appendix**. Source data and genetic files are available in DRYAD (https://doi.org/10.5061/dryad.zgmsbccmz). For review purposes, source data can be accessed via a temporary link (https://rb.gy/lpdr3a).

#### Supplementary materials and methods

**1.** Transmembrane domain removal and codon optimization of cytochromes P450 enzymes

**2.** Algae cultivation, plasmid design, transformation, and screening

**3.** Gas chromatography-flame ionization detection/mass spectrometry GC–FID/MS method

#### SI Appendix Figures

**Fig. S1.** Fluorescence-based screening of *C. reinhardtii* transformants expressing plasmids A.

**Fig. S2.** Fluorescence-based screening of *C. reinhardtii* transformants expressing plasmids B.

**Fig. S3.** Fluorescence-based screening of *C. reinhardtii* transformants expressing plasmids C.

**Fig. S4.** Fluorescence-based screening of *C. reinhardtii* transformants expressing CYP plasmids.

**Fig. S5.** Calibration curves for sesquiterpenoid standards

**Fig. S6.** Relative abundance of sesquiterpenoids produced in *C. reinhardtii* expressing different CYPs

**Fig. S7.** Functionalization efficiency of tested CYPs

#### SI Appendix Tables

**Table. S1.** List of plasmids used in this study.

**Table. S2.** Quantification of sesquiterpenoids from synthases localized in different subcellular compartments

**Table S3.** Two-way ANOVA sesquiterpenoids from synthases localized in different subcellular compartments.

**Table. S4.** Peak areas (GC-FID) *C. reinhardtii* strains harboring STPS and CYPs

**Table. S5.** Conversion efficiency (%) of CYPs

**Table. S6.** Extraction capacities of different fluorinate solvents

**Table. S7.** GC-MS of standard terpenoid mixture.

**Table. S8.** GC-MS results of engineered *C. reinhardtii* strains

**Table. S9.** Mass spectra of sesquiterpenoids identified by GC – MS.

#### SI Appendix Files

**File S1.** Genetic constructs used in this study.

**File S2.** GC-MS/FID chromatograms for *C. reinhardtii* strains harboring plasmids A.

**File S3.** GC-MS/FID chromatograms for *C. reinhardtii* strains harboring plasmids B.

**File S4.** GC-MS/FID chromatograms for *C. reinhardtii* strains harboring plasmids C.

**File S5.** GC-MS/FID chromatograms for *C. reinhardtii* strains co-expressing plasmids B and CYP.

**File S6.** GC-MS/FID chromatograms for *C. reinhardtii* strains co-expressing plasmids B and CYP grown with different carbon sources.

**File S7.** Raw GC-MS/FID data.

### Supplementary materials and methods

#### 1. Transmembrane domain removal and codon optimization of cytochromes P450 enzymes

To enable soluble expression in the plastid stroma, transmembrane (TM) domains were removed from all cytochrome P450 (CYP) amino acid sequences. TM domains were initially predicted using the TMHMM - 2.0 server (http://www.cbs.dtu.dk/services/TMHMM/), which employs hidden Markov models to identify probable membrane-spanning regions. AlphaFold models were then generated for each full-length CYP sequence and visually inspected to confirm the presence of low-structured N-terminal regions corresponding to TM domains. Based on these analyses, N-terminal TM domains were identified and removed from each CYP sequence, with the care of preserving essential residues immediately following the TM domain. The modified sequences were re-analyzed using TMHMM - 2.0 and AlphaFold to verify the absence of TM domains and the expected soluble protein structure. For optimal expression in *C. reinhardtii*, the modified amino acid sequences were back-translated and codon-optimized using the Intronserter web tool (https://bibiserv.cebitec.uni-bielefeld.de/intronserter). This tool implements back translation and codon optimization based on C. reinhardtii-specific codon usage tables, removes undesired sequence elements, and performs systematic insertion of the first intron of C. reinhardtii ribulose-1,5-bisphosphate carboxylase/oxygenase small subunit 2 (rbcS2i1) to minimize exon lengths. The resulting nucleotide sequences, optimized for codon usage and intron content, were synthesized de novo and subcloned by Genscript (Piscataway, NJ, USA).

#### 2. Algae cultivation, plasmid design, transformation, and screening

Experiments employed a genetically engineered *C. reinhardtii* strain derived from UPN22 (a UVM4 derivative) optimized for enhanced terpenoid biosynthesis. Engineered strains were cultured in TAPhi-$NO_3$ liquid medium in microtiter plates with agitation or on solid agar under LED illumination at 150 µmol $m^{-2}$ $s^{-1}$. *C. reinhardtii* nuclear genome transformation used linearized plasmid DNA (XbaI + KpnI restriction enzymes, Thermo Scientific FastDigest) via a glass-bead protocol. Each transformation utilized 10 µg of DNA. After an 8-hour recovery in liquid TAPhi-$NO_3$ medium under low light, algal cells were plated on medium containing antibiotics: paromomycin (10 µg $mL^{-1}$), spectinomycin (200 µg $mL^{-1}$), hygromycin B (15 µg $mL^{-1}$), or zeocin (15 µg $mL^{-1}$), singly or in combinations for desired selection. Plates were exposed to continuous light for 7 days before colony selection. A PIXL robot (Singer Instruments, Watchet, UK) transferred up to 384 colonies per transformation to TAPhi-NO3 agar plates. After 3 days, a ROTOR robot replicated colonies onto plates containing amido black (150 µg mL–1) for fluorescence screening. All algae-optimized constructs were expressed as fusion proteins with mVenus (yellow fluorescent protein), mKOkappa (orange fluorescent protein), or monomeric teal fluorescent protein 1 (mTFP1). Transgene expression was evaluated by fluorescence imaging of colonies on agar plates. Fluorescence screening protocol: **1)** Chlorophyll fluorescence (colony presence): 3-second excitation at 475/20 nm, emission at 640/160 nm. **2)** Teal (mTFP1) fluorescence: 2.5-minute exposure, 420/20 nm excitation emission 480/20 nm. **3)** Yellow (mVenus) fluorescence: 30-second exposure, excitation 504/10 nm, emission 530/10 nm. **4)** Orange (mKOk) fluorescence: 30-second exposure, excitation 504/10 nm, emission 600/10 nm. Colonies exhibiting strong fluorescent-protein signals were selected from 96-colony arrays, transferred to 12-well plates with 2 mL of liquid TAPhi-NO3 medium, and cultured with agitation at 160 rpm.

#### 3. Gas chromatography-flame ionization detection/mass spectrometry methods

Analysis was performed using an Agilent 7890A gas chromatograph with a mass spectrometer and flame-ionization detector (GC-MS/FID). The system included a 5975C inert MSD with a triple-axis detector and a DB-5MS column (30 m × 0.25 mm i.d., 0.25 μm film thickness). Temperatures were set to 250 °C for the injector and interface and 220 °C for the ion source. An autosampler (G4513A, Agilent) injected 1 μL of sample in splitless mode. Helium carrier gas flow was maintained at 1 mL $min^{-1}$. The GC oven temperature program was as follows: 80 °C for 1 min, increased to 120 °C at 10 °C $min^{-1}$, to 160 °C at 3 °C $min^{-1}$, and to 280 °C at 10 °C $min^{-1}$, with a final hold for 3 min. Mass spectra were recorded after a 14-minute solvent delay, scanning 50–350 m/z at 50 scans per second. Chromatograms were analyzed using MassHunter Workstation software version B.08.00 (Agilent). Terpenoids were identified using the National Institute of Standards and Technology (NIST) library (Gaithersburg, MD, USA). Further identification utilized purified standard calibration curves (1–1200 μM in dodecane) for δ-guaiene (CAT#B942760), patchoulol (CAT#P206200), santalene (CAT#S15065), valerianol (CAT#V914000, Toronto Research Chemicals, ON, Canada), bisabolol (CAT#95426), valencene (CAT#06808), and cedrene (CAT#22133, Sigma-Aldrich, MO, USA). For compound identification, retention-time acquisition, internal digital library calibration, and method development, we used a set of 12 microampules containing a standard terpene mixture of 98 terpenes at 1 mM in methanol (CAT# MSITPN101, MetaSci, Canada).

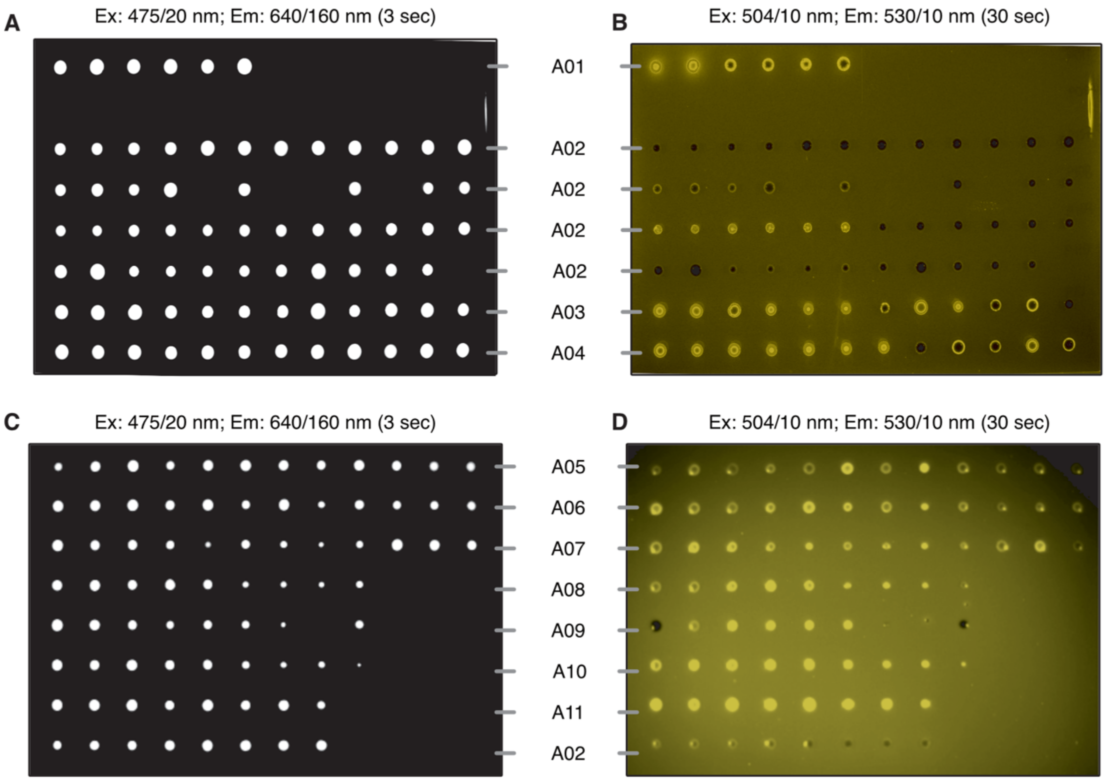


**Fig. S1.** Fluorescence-based screening of *C. reinhardtii* transformants expressing plasmids A. **(A, C)** Chlorophyll autofluorescence (Ex: 475/20 nm; Em: 640/160 nm, 3 sec exposure) indicating colony presence. **(B, D)** Yellow fluorescent protein (mVenus) signal (Ex: 504/10 nm; Em: 530/10 nm, 30-sec exposure) showing transgene expression. Rows labeled A01-A11 represent different transformed lines.

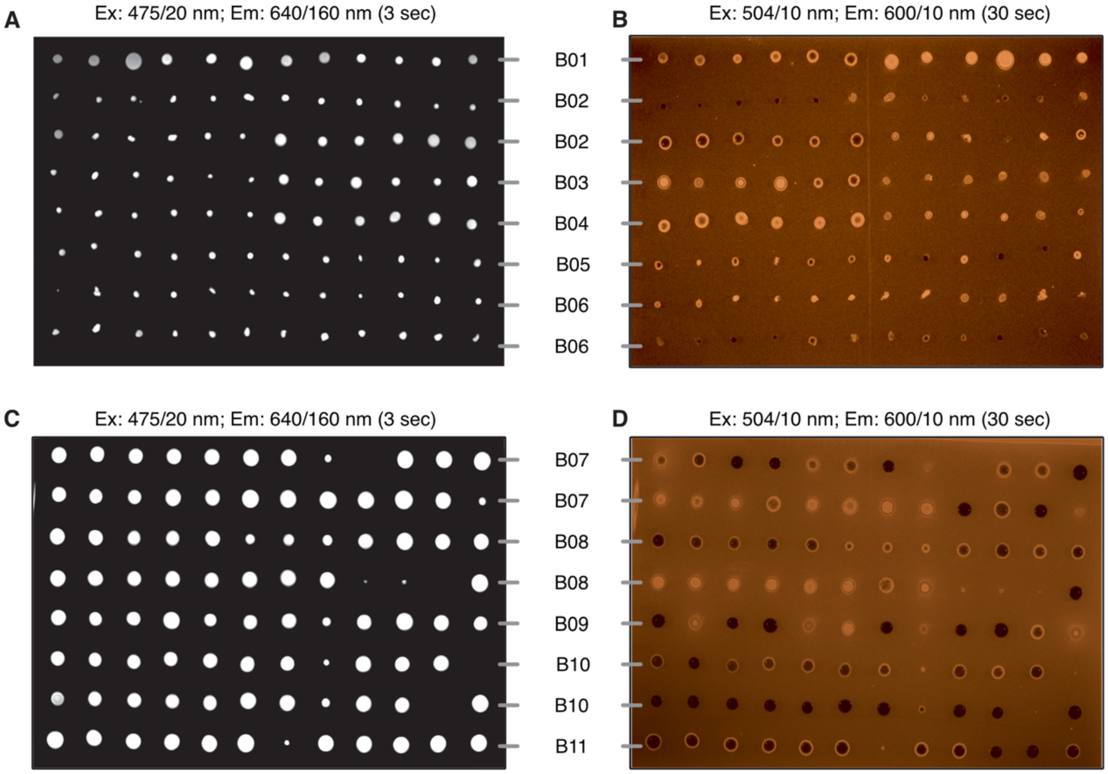


Fig. S2. Fluorescence-based screening of *C. reinhardtii* transformants expressing plasmids B. (A, C) Chlorophyll autofluorescence (Ex: 475/20 nm; Em: 640/160 nm, 3 sec exposure) indicating colony presence. (B, D) Orange fluorescent protein (mKOkappa) signal (Ex: 504/10 nm; Em: 600/10 nm, 30-sec exposure) showing transgene expression. Rows labeled B01-B11 represent different transformed lines.

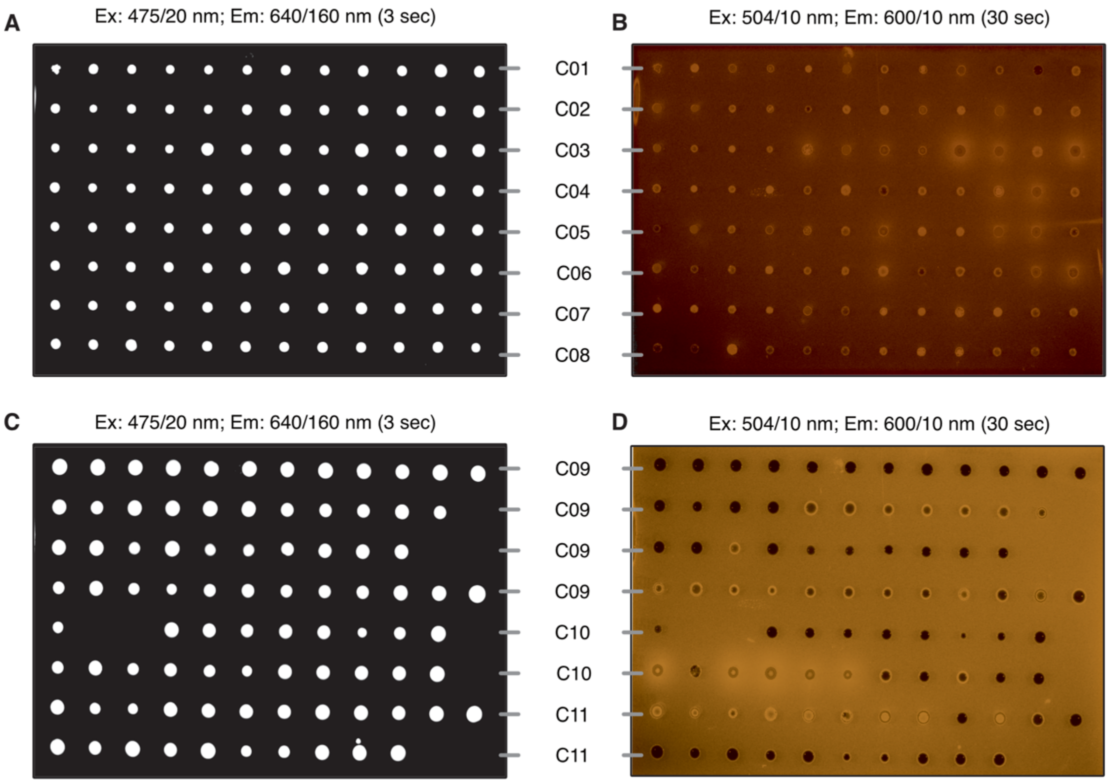


**Fig. S3.** Fluorescence-based screening of *C. reinhardtii* transformants expressing plasmids C. **(A, C)** Chlorophyll autofluorescence (Ex: 475/20 nm; Em: 640/160 nm, 3 sec exposure) indicating colony presence. **(B, D)** Orange fluorescent protein (mKOkappa) signal (Ex: 504/10 nm; Em: 600/10 nm, 30 sec exposure) showing transgene expression. Rows labeled C01-C11 represent different transformed lines.

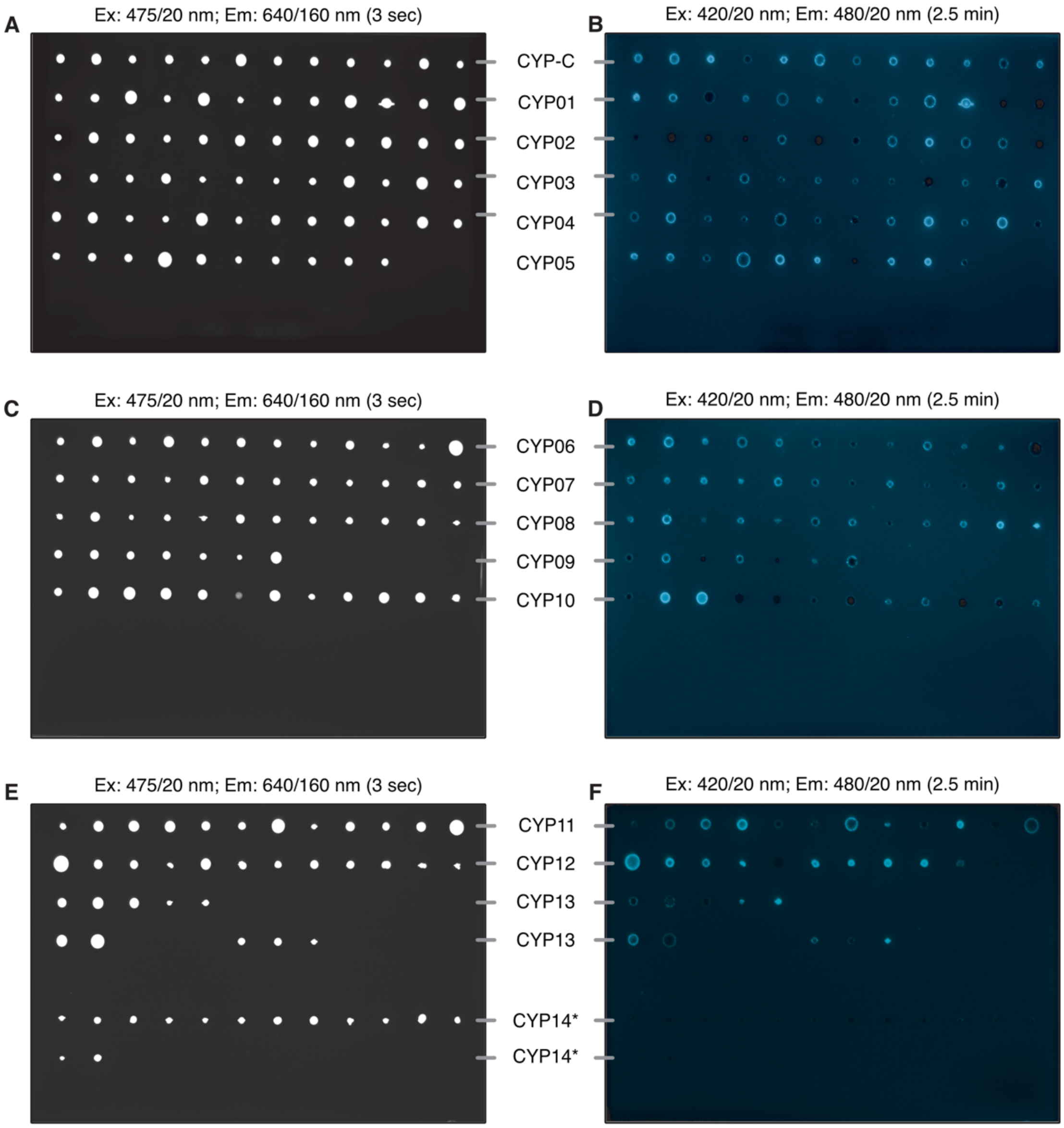


**Fig. S4.** Fluorescence-based screening of **C. reinhardtii** transformants expressing cytochrome P450 (CYP) plasmids.

**(A, C, E)** Chlorophyll autofluorescence (Ex: 475/20 nm; Em: 640/160 nm, 3 sec exposure) indicating colony presence. **(B, D, F)** Teal fluorescent protein (mTFP1) signal (Ex: 420/20 nm; Em: 480/20 nm, 2.5 min exposure) showing transgene expression. Rows labeled CYP-C to CYP14 represent different CYP-expressing lines.

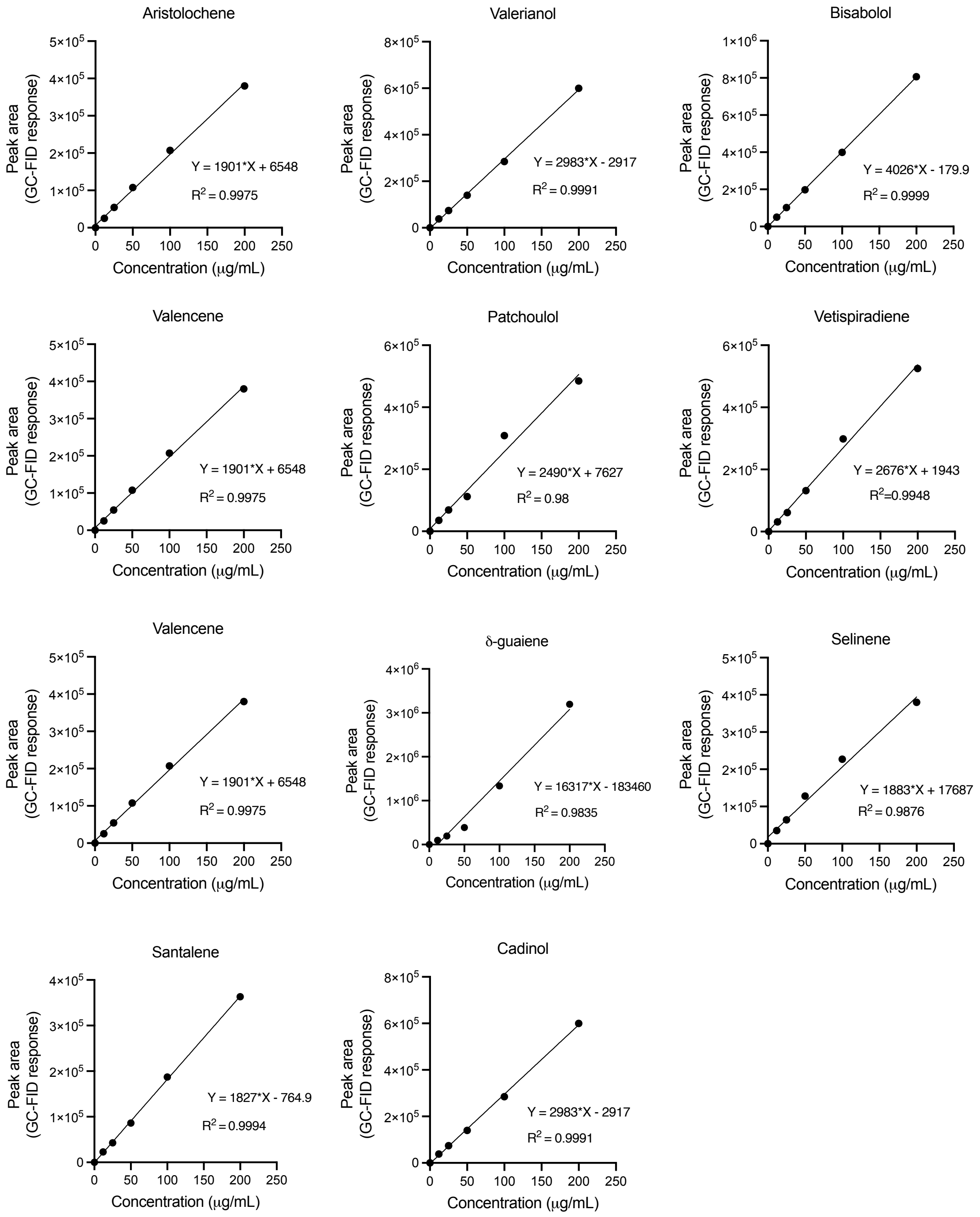


**Fig. S5.** Calibration curves for sesquiterpenoid standards.

Standard curves used for quantification of sesquiterpenoids produced in *C. reinhardtii.*

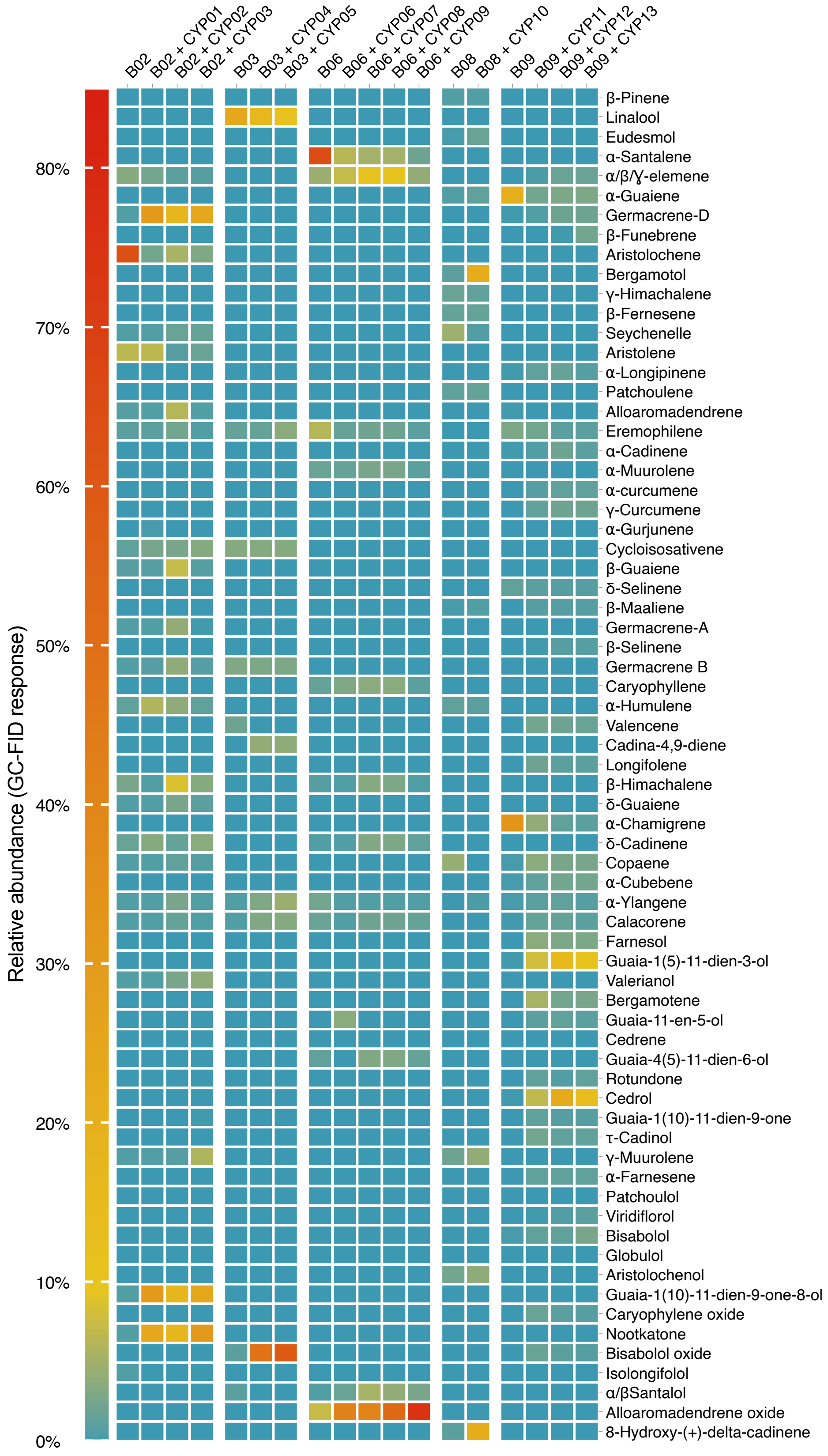
B02
B02 + CYP01
B02 + CYP02
B02 + CYP03
B03
B03 + CYP04
B03 + CYP05
B06
B06 + CYP06
B06 + CYP07
B06 + CYP08
B06 + CYP09
B08
B08 + CYP10
B09
B09 + CYP11
B09 + CYP12
B09 + CYP13
Relative abundance (GC-FID response)
80%
70%
60%
50%
40%
30%
20%
10%
0%
β-Pinene
Linalool
Eudesmol
α-Santalene
α/β/Ɣ-elemene
α-Guaiene
Germacrene-D
β-Funebrene
Aristolochene
Bergamotol
γ-Himachalene
β-Fernesene
Seychenelle
Aristolene
α-Longipinene
Patchoulene
Alloaromadendrene
Eremophilene
α-Cadinene
α-Muurolene
α-curcumene
γ-Curcumene
α-Gurjunene
Cycloisosativene
β-Guaiene
δ-Selinene
β-Maaliene
Germacrene-A
β-Selinene
Germacrene B
Caryophyllene
α-Humulene
Valencene
Cadina-4,9-diene
Longifolene
β-Himachalene
δ-Guaiene
α-Chamigrene
δ-Cadinene
Copaene
α-Cubebene
α-Ylangene
Calacorene
Farnesol
Guaia-1(5)-11-dien-3-ol
Valerianol
Bergamotene
Guaia-11-en-5-ol
Cedrene
Guaia-4(5)-11-dien-6-ol
Rotundone
Cedrol
Guaia-1(10)-11-dien-9-one
τ-Cadinol
γ-Muurolene
α-Farnesene
Patchoulol
Viridiflorol
Bisabolol
Globulol
Aristolochenol
Guaia-1(10)-11-dien-9-one-8-ol
Caryophylene oxide
Nootkatone
Bisabolol oxide
Isolongifolol
α/βSantalol
Alloaromadendrene oxide
8-Hydroxy-(+)-delta-cadinene

**Fig. S6.** Relative abundance of sesquiterpenoids produced in *C. reinhardtii* expressing different CYPs. Comparison of sesquiterpenoid profiles among strains expressing various CYPs.

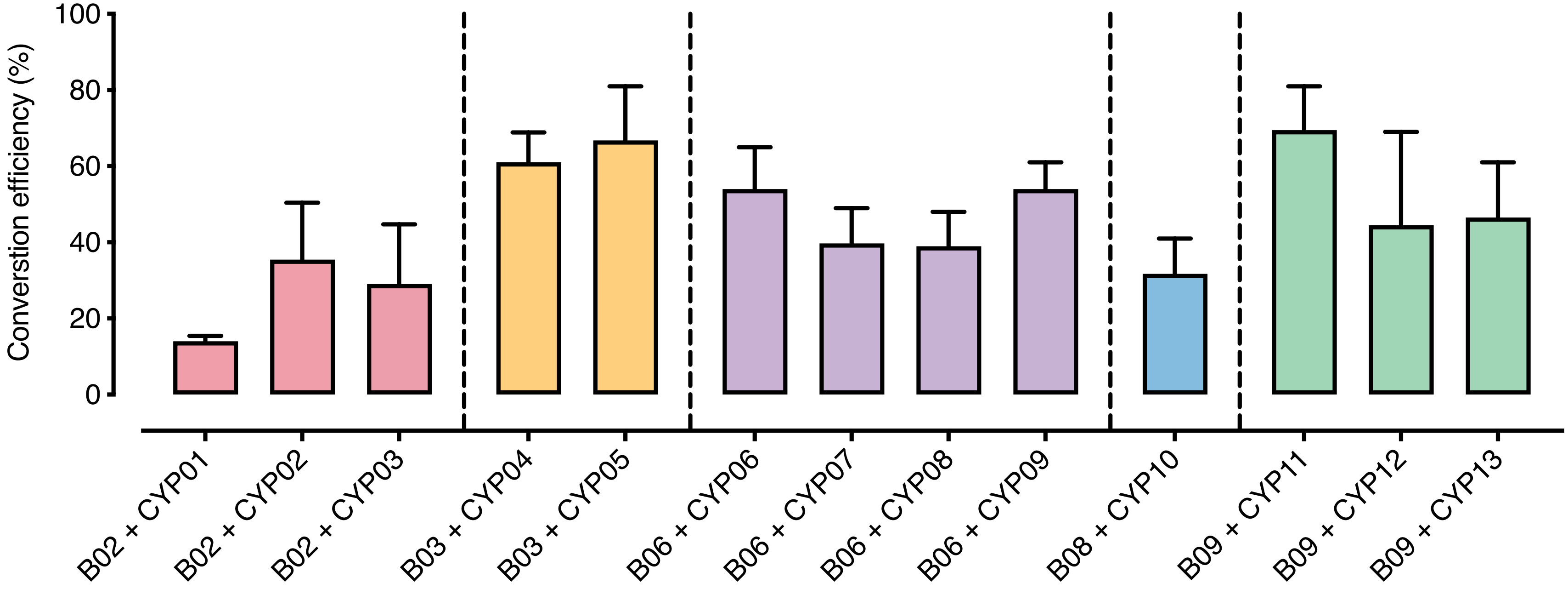


**Fig. S7.** Functionalization efficiency of tested cytochrome P450 enzymes (CYPs).
Percentage of substrate converted to functionalized products by each CYP.

**Table S1.** Plasmids used in this study.

| Plasmid | ID | Gene name | Target peptide | Origin | UniProt | Selection | Reporter | Comment |
|---|---|---|---|---|---|---|---|---|
| 1 | - | SQS amiRNA | CAH1 | *C. reinhardtii* | - | Spectinomycin | *g*Luc | SQS.k.d |
| 2 | - | *Cr*BKT | CTP | *C. reinhardtii* | - | Spectinomycin | N/A | - |
| 3 | A01 | mVenus | N/A | *Aequorea victoria* | - | Paromomycin | mVenus | YFP |
| 4 | B01 | mKOkappa, *Sc*Erg20 | CTP | *Verrillofungia concinna, Saccharomyces cerevisiae* | - | Hygromycin | mKOκ | Orange FP |
| 5 | C01 | mKOkappa, *Ec*ispA | CTP | *V. concinna, Escherichia coli* | - | Hygromycin | mKOκ | Orange FP |
| 6 | CYP-C | mTFP1 | CTP | *Clavularia spp.* | - | Bleomycin | mTFP1 | Teal FP |
| 7 | A02 | PcAS | N/A | *Penicillium chrysogenum* | A0A167QKS4 | Paromomycin | mVenus | Aristolochene |
| 8 | B02 | *Pc*AS, *Sc*Erg20 | CTP | *P. chrysogenum, S.cerevisiae* | A0A167QKS4, P08524 | Hygromycin | mKOκ | Aristolochene |
| 9 | C02 | *Pc*AS, *EcispA* | CTP | *P. chrysogenum, E. coli* | A0A167QKS4, P22939 | Hygromycin | mKOκ | Aristolochene |
| 10 | CYP01 | SLS1 | CTP | *Catharanthus roseus* | Q05047 | Bleomycin | mTFP1 | Aristolochene |
| 11 | CYP02 | prx4 | CTP | *Penicillium roqueforti* (strain FM164) | W6QP06 | Bleomycin | mTFP1 | Aristolochene |
| 12 | CYP03 | ORF5 | CTP | *P. roqueforti* (strain FM164) | W6Q3Z9 | Bleomycin | mTFP1 | Aristolochene |
| 13 | A03 | *Cn*VS | N/A | *Callitropsis nootkatensis* | A0A167QKS4 | Paromomycin | mVenus | Valencene |

Table S1. Cont.

| Plasmid | ID | Gene name | Target peptide | Origin | UniProt | Selection | Reporter | Comment |
|---|---|---|---|---|---|---|---|---|
| 14 | B03 | *Cn*VS | CTP | *C. nootkatensis, S. cerevisiae* | A0A167QKS4, P08524 | Hygromycin | mKOκ | Valencene |
| 15 | C03 | *Cn*VS | CTP | *C. nootkatensis, E. coli* | A0A167QKS4, P22939 | Hygromycin | mKOκ | Valencene |
| 16 | CYP04 | C*n*VO | CTP | *C. nootkatensis* | V9N2L5 | Bleomycin | mTFP1 | Valencene |
| 17 | CYP05 | CYP71AV8 | CTP | *Cichorium intybus* | E1B2Z9 | Bleomycin | mTFP1 | Valencene |
| 18 | A04 | *Ag*SS | N/A | *Abies grandis* | O64404 | Paromomycin | mVenus | Selinene |
| 19 | B04 | *Ag*SS, *Sc*Erg20 | CTP | *A. grandis, S. cerevisiae* | O64404, P08524 | Hygromycin | mKOκ | Selinene |
| 20 | C04 | *Ag*SS, *Ec*ispA | CTP | *A. grandis, E.coli* | O64404, P22939 | Hygromycin | mKOκ | Selinene |
| 21 | A05 | *As*VS | N/A | *Aquilaria Sinensis* | A0A411G8M5 | Paromomycin | mVenus | Vetispiradiene |
| 22 | B05 | *As*VS, *Sc*Erg20 | CTP | *A. Sinensis, S. cerevisiae* | A0A411G8M5, P08524 | Hygromycin | mKOκ | Vetispiradiene |
| 23 | C05 | *As*VS, *Ec*ispA | CTP | *A. Sinensis, E. coli* | A0A411G8M5, P22939 | Hygromycin | mKOκ | Vetispiradiene |
| 24 | A06 | *Cl*SS | N/A | *Clausena lansium* | E5LLI1 | Paromomycin | mVenus | Santalene |
| 25 | B06 | *Cl*SS, ScErg20 | CTP | *C. lansium, S. cerevisiae* | E5LLI1, P08524 | Hygromycin | mKOκ | Santalene |

Table S1. Cont.

| Plasmid | ID | Gene name | Target peptide | Origin | UniProt | Selection | Reporter | Comment |
|---|---|---|---|---|---|---|---|---|
| 26 | C06 | *Cl*SS, EcispA | CTP | *C. lansium, E. coli* | E5LLI1, P22939 | Hygromycin | mKOκ | Santalene |
| 27 | CYP06 | CYP736A167 | CTP | *S. album* | A0A142I6X1 | Bleomycin | mTFP1 | Santalene |
| 28 | CYP07 | CYP76F39 | CTP | *S. album* | V5RH69 | Bleomycin | mTFP1 | Santalene |
| 29 | CYP08 | CYP76F39 | CTP | *S. album* | V5RGD0 | Bleomycin | mTFP1 | Santalene |
| 30 | CYP09 | CYP76F40 | CTP | *S. album* | V5REU4 | Bleomycin | mTFP1 | Santalene |
| 31 | A07 | *Cc*BS | N/A | *Cynara cardunculus* | A0A103XRG2 | Paromomycin | mVenus | Bisabolol |
| 32 | B07 | *Cc*BS, *ScErg20* | CTP | *C. cardunculus, S. cerevisiae* | A0A103XRG2, P08524 | Hygromycin | mKOκ | Bisabolol |
| 33 | C07 | *Cc*BS, *EcispA* | CTP | *C. cardunculus, E. coli* | A0A103XRG2, P22939 | Hygromycin | mKOκ | Bisabolol |
| 34 | A08 | *La*CS | N/A | *Lavandula angustifolia* | U3LW50 | Paromomycin | mVenus | Cadinol |
| 35 | B08 | *La*CS, *ScErg20* | CTP | *L. angustifolia, S. cerevisiae* | U3LW50, P08524 | Hygromycin | mKOκ | Cadinol |
| 36 | C08 | *La*CS, *EcispA* | CTP | *L. angustifolia, E. coli* | U3LW50, P22939 | Hygromycin | mKOκ | Cadinol |
| 37 | CYP10 | CAD8H-1 | CTP | *Gossypium hirsutum* | A0A0N9H930 | Bleomycin | mTFP1 | Cadinol |
| 38 | A09 | *AsGS* | N/A | *A.sinensis* | K9MPP8 | Paromomycin | mVenus | Guaiene |

**Table S1**. Cont.

| Plasmid | ID | Gene name | Target peptide | Origin | UniProt | Selection | Reporter | Comment |
|---|---|---|---|---|---|---|---|---|
| 39 | B09 | *AsGS, ScErg20* | CTP | *A.sinensis, S. cerevisiae* | K9MPP8, P08524 | Hygromycin | mKOκ | Guaiene |
| 40 | C09 | *AsGS, EcispA* | CTP | *A. sinensis, E. coli* | K9MPP8, P22939 | Hygromycin | mKOκ | Guaiene |
| 41 | CYP11 | CYP71BE5 | CTP | *Vitis vinifera (Grape)* | F6I534 | Bleomycin | mTFP1 | Guaiene |
| 42 | CYP12 | CYP71BA1 | CTP | *Zingiber zerumbet* | E3W9C4 | Bleomycin | mTFP1 | Guaiene |
| 43 | CYP13 | CYP71D55 | CTP | *Hyoscyamus muticus* | A6YIH8 | Bleomycin | mTFP1 | Guaiene |
| 44 | A10 | *Ch*VOS | N/A | *Camellia hiemalis* | A0A348AUV5 | Paromomycin | mVenus | Valerianol |
| 45 | B10 | *Ch*VOS, *ScErg20* | CTP | *C.hiemalis, S. cerevisiae* | A0A348AUV5, P08524 | Hygromycin | mKOκ | Valerianol |
| 46 | C10 | *Ch*VOS, *EcispA* | CTP | *C. hiemalis, E. coli* | A0A348AUV5, P22939 | Hygromycin | mKOκ | Valerianol |
| 47 | A11 | *Pc*PS2 | N/A | *Pogostemon cablin* | Q49SP3* | Paromomycin | mVenus | Patchoulol |
| 48 | B11 | *Pc*PS2, *ScErg20* | CTP | *P. cablin* | Q49SP3*, P08524 | Hygromycin | mKOκ | Patchoulol |
| 49 | C11 | *Pc*PS2, *EcispA* | CTP | *P.cablin* | Q49SP3*, P22939 | Hygromycin | mKOκ | Patchoulol |
| 50 | CYP14* | ORF6 | CTP | *Penicillium roqueforti* (strain FM164) | W6QB15 | Bleomycin | mTFP1 | Aristolochene |

**Table S2.** Quantification of sesquiterpenoids from synthases localized in different subcellular compartments

| Sesquiterpenoid | Cytoplasm (A) (μg/L) | | Chloroplast (B) (μg/L) | | Chloroplast (C) (μg/L) | |
|---|---|---|---|---|---|---|
| | Mean | SD | Mean | SD | Mean | SD |
| Blank | 0 | 0 | 0 | 0 | 0 | 0 |
| Aristolochene | 355 | 107 | 165 | 91 | 196 | 208 |
| Valencene | 338 | 56 | 215 | 120 | 315 | 160 |
| Selinene | 352 | 146 | 253 | 100 | 290 | 125 |
| Vetispiradiene | 321 | 82 | 175 | 98 | 194 | 131 |
| Santalene | 365 | 76 | 324 | 135 | 245 | 167 |
| Bisabolol | 1172 | 130 | 1024 | 187 | 135 | 84 |
| Cadinol | 1199 | 234 | 977 | 305 | 406 | 208 |
| Guaiene | 1707 | 296 | 893 | 149 | 1294 | 186 |
| Valerianol | 1563 | 83 | 587 | 218 | 954 | 354 |
| Patchoulol | 2004 | 432 | 1562 | 187 | 1557 | 541 |

**Table S3.** Two-way ANOVA sesquiterpenoids from synthases localized in different subcellular compartments

| Sesquiterpenoid | Turkey's multiple compartison | Mean 1 | Mean 2 | Mean Diff. | 95.00% CI of diff. | Adjusted P Value |
|---|---|---|---|---|---|---|
| Negative | Cytoplasm vs. Chloroplast (Erg20) | 0 | 0 | 0 | -337.4 to 337.4 | >0.9999 |
| | Cytoplasm vs. Chloroplast (ispA) | 0 | 0 | 0 | -337.4 to 337.4 | >0.9999 |
| | Chloroplast (Erg20) vs. Chloroplast (ispA) | 0 | 0 | 0 | -337.4 to 337.4 | >0.9999 |
| Aristolochene | Cytoplasm vs. Chloroplast (Erg20) | 355.3 | 164.5 | 190.8 | -146.5 to 528.2 | 0.3734 |
| | Cytoplasm vs. Chloroplast (ispA) | 355.3 | 195.5 | 159.8 | -177.5 to 497.2 | 0.4996 |
| | Chloroplast (Erg20) vs. Chloroplast (ispA) | 164.5 | 195.5 | -30.99 | -368.3 to 306.4 | 0.974 |
| Valencene | Cytoplasm vs. Chloroplast (Erg20) | 338.1 | 215.4 | 122.8 | -214.6 to 460.1 | 0.663 |
| | Cytoplasm vs. Chloroplast (ispA) | 338.1 | 315 | 23.11 | -314.2 to 360.5 | 0.9855 |
| | Chloroplast (Erg20) vs. Chloroplast (ispA) | 215.4 | 315 | -99.65 | -437.0 to 237.7 | 0.7624 |
| Selinene | Cytoplasm vs. Chloroplast (Erg20) | 352 | 253.3 | 98.75 | -238.6 to 436.1 | 0.7661 |
| | Cytoplasm vs. Chloroplast (ispA) | 352 | 290.3 | 61.75 | -275.6 to 399.1 | 0.9008 |
| | Chloroplast (Erg20) vs. Chloroplast (ispA) | 253.3 | 290.3 | -37 | -374.4 to 300.4 | 0.9632 |
| Vetispiradiene | Cytoplasm vs. Chloroplast (Erg20) | 320.8 | 175 | 145.8 | -191.6 to 483.1 | 0.561 |
| | Cytoplasm vs. Chloroplast (ispA) | 320.8 | 193.8 | 127 | -210.4 to 464.4 | 0.6442 |
| | Chloroplast (Erg20) vs. Chloroplast (ispA) | 175 | 193.8 | -18.75 | -356.1 to 318.6 | 0.9904 |

Table S3. Cont.

| Sesquiterpenoid | Turkey's multiple compartison | Mean 1 | Mean 2 | Mean Diff. | 95.00% CI of diff. | Adjusted P Value |
|---|---|---|---|---|---|---|
| Santalene | Cytoplasm vs. Chloroplast (Erg20) | 365.3 | 323.7 | 41.56 | -295.8 to 378.9 | 0.9538 |
| | Cytoplasm vs. Chloroplast (ispA) | 365.3 | 244.6 | 120.7 | -216.7 to 458.0 | 0.6722 |
| | Chloroplast (Erg20) vs. Chloroplast (ispA) | 323.7 | 244.6 | 79.12 | -258.2 to 416.5 | 0.8426 |
| Bisabolol | Cytoplasm vs. Chloroplast (Erg20) | 1172 | 1024 | 147.6 | -189.8 to 484.9 | 0.5529 |
| | Cytoplasm vs. Chloroplast (ispA) | 1172 | 134.5 | 1038 | 700.2 to 1375 | <0.0001 |
| | Chloroplast (Erg20) vs. Chloroplast (ispA) | 1024 | 134.5 | 889.9 | 552.6 to 1227 | <0.0001 |
| Cadinol | Cytoplasm vs. Chloroplast (Erg20) | 1199 | 976.6 | 222.2 | -115.2 to 559.5 | 0.2647 |
| | Cytoplasm vs. Chloroplast (ispA) | 1199 | 405.8 | 793 | 455.7 to 1130 | <0.0001 |
| | Chloroplast (Erg20) vs. Chloroplast (ispA) | 976.6 | 405.8 | 570.8 | 233.5 to 908.2 | 0.0003 |
| Guaiene | Cytoplasm vs. Chloroplast (Erg20) | 1707 | 893 | 814.3 | 476.9 to 1152 | <0.0001 |
| | Cytoplasm vs. Chloroplast (ispA) | 1707 | 1294 | 413.2 | 75.80 to 750.5 | 0.0122 |
| | Chloroplast (Erg20) vs. Chloroplast (ispA) | 893 | 1294 | -401.1 | -738.5 to -63.75 | 0.0155 |
| Valerianol | Cytoplasm vs. Chloroplast (Erg20) | 1563 | 587.4 | 975.6 | 638.2 to 1313 | <0.0001 |
| | Cytoplasm vs. Chloroplast (ispA) | 1563 | 953.5 | 609.5 | 272.1 to 946.8 | 0.0001 |
| | Chloroplast (Erg20) vs. Chloroplast (ispA) | 587.4 | 953.5 | -366.1 | -703.5 to -28.75 | 0.0301 |
| Patchoulol | Cytoplasm vs. Chloroplast (Erg20) | 2004 | 1562 | 442.6 | 105.2 to 780.0 | 0.0066 |
| | Cytoplasm vs. Chloroplast (ispA) | 2004 | 1557 | 447.3 | 110.0 to 784.7 | 0.006 |
| | Chloroplast (Erg20) vs. Chloroplast (ispA) | 1562 | 1557 | 4.737 | -332.6 to 342.1 | 0.9994 |

**Table S4.** Peak areas (GC-FID) of STPs from *C. reinhardtii* strains expressing STPS and CYPs

| Strain | Sesquiterpenoid (Peak area) | | | | Functionalized sesquiterpenoid (Peak area) | | | |
|---|---|---|---|---|---|---|---|---|
| B01 | 0.0E+00 | 0.0E+00 | 0.0E+00 | 0.0E+00 | 0.0E+00 | 0.0E+00 | 0.0E+00 | 0.0E+00 |
| B02 | 3.8E+04 | 4.5E+04 | 4.2E+04 | 4.2E+04 | 5.6E+03 | 6.7E+03 | 6.3E+03 | 6.3E+03 |
| B02 + CYP01 | 4.9E+02 | 3.2E+02 | 3.3E+02 | 3.3E+02 | 4.9E+04 | 3.2E+04 | 3.3E+04 | 3.3E+04 |
| B02 + CYP02 | 3.2E+02 | 4.2E+02 | 4.8E+02 | 2.7E+02 | 3.2E+04 | 4.2E+04 | 4.8E+04 | 2.7E+04 |
| B02 + CYP03 | 1.6E+02 | 3.2E+02 | 4.8E+02 | 3.2E+02 | 1.6E+04 | 3.2E+04 | 4.8E+04 | 3.2E+04 |
| B03 | 8.3E+03 | 7.8E+03 | 1.0E+04 | 1.4E+04 | 1.1E+03 | 1.0E+03 | 1.3E+03 | 1.8E+03 |
| B03 + CYP04 | 2.1E+03 | 2.1E+03 | 2.0E+03 | 3.3E+03 | 2.1E+05 | 2.1E+05 | 2.0E+05 | 3.3E+05 |
| B03 + CYP05 | 2.0E+03 | 2.1E+03 | 2.4E+03 | 2.0E+03 | 2.0E+05 | 2.1E+05 | 2.4E+05 | 2.0E+05 |
| B06 | 2.1E+04 | 1.9E+04 | 2.3E+04 | 2.5E+04 | 1.5E+02 | 1.1E+02 | 1.2E+02 | 1.5E+02 |
| B06 + CYP06 | 1.5E+02 | 1.2E+02 | 1.3E+02 | 1.5E+02 | 2.1E+04 | 1.9E+04 | 2.3E+04 | 2.5E+04 |
| B06 + CYP07 | 5.3E+02 | 6.9E+02 | 6.9E+02 | 1.1E+03 | 2.0E+04 | 1.8E+04 | 2.2E+04 | 2.4E+04 |
| B06 + CYP08 | 3.9E+02 | 4.8E+02 | 8.4E+02 | 1.0E+03 | 2.0E+04 | 1.8E+04 | 2.2E+04 | 2.4E+04 |
| B06 + CYP09 | 2.9E+03 | 2.0E+03 | 1.6E+03 | 1.2E+03 | 1.8E+04 | 1.7E+04 | 2.1E+04 | 2.4E+04 |
| B08 | 7.4E+04 | 3.6E+04 | 5.2E+04 | 3.5E+04 | 3.7E+03 | 1.8E+03 | 2.6E+03 | 1.7E+03 |
| B08 + CYP09 | 5.3E+03 | 3.1E+03 | 3.6E+03 | 2.4E+03 | 5.3E+05 | 3.1E+05 | 3.6E+05 | 2.4E+05 |
| B09 | 5.6E+04 | 5.0E+04 | 6.6E+04 | 5.7E+04 | 9.0E+03 | 8.1E+03 | 1.1E+04 | 9.1E+03 |
| B09 + CYP11 | 7.0E+03 | 7.3E+03 | 7.0E+03 | 7.6E+03 | 7.0E+05 | 7.3E+05 | 7.0E+05 | 7.6E+05 |
| B09 + CYP12 | 8.0E+03 | 7.0E+03 | 7.4E+03 | 7.4E+03 | 8.0E+05 | 7.0E+05 | 7.4E+05 | 7.4E+05 |
| B09 + CYP13 | 7.4E+03 | 5.7E+03 | 7.2E+03 | 7.1E+03 | 7.4E+05 | 5.7E+05 | 7.2E+05 | 7.1E+05 |

**Table S5.** Conversion efficiency (%) of CYPs

| **Strain (CYP)** | **Conversation efficiency (%)** | | | | | |
|---|---|---|---|---|---|---|
| | R1 | R2 | R3 | R4 | Mean | SD |
| B02 + CYP01 | 58 | 61 | 44 | 53 | 54.0 | 7.4 |
| B02 + CYP02 | 32 | 19 | 55 | 36 | 35.5 | 14.9 |
| B02 + CYP03 | 21 | 15 | 29 | 51 | 29.0 | 15.7 |
| B03 + CYP04 | 62 | 71 | 52 | 59 | 61.0 | 7.9 |
| B03 + CYP05 | 68 | 81 | 55 | 63 | 66.8 | 10.9 |
| B06 + CYP06 | 65 | 52 | 58 | 41 | 54.0 | 10.2 |
| B06 + CYP07 | 65 | 52 | 58 | 41 | 54.0 | 10.2 |
| B06 + CYP08 | 42 | 49 | 32 | 36 | 39.8 | 7.4 |
| B06 + CYP09 | 34 | 39 | 35 | 48 | 39.0 | 6.4 |
| B08 + CYP10 | 21 | 36 | 29 | 41 | 31.8 | 8.7 |
| B09 + CYP11 | 81 | 65 | 71 | 61 | 69.5 | 8.7 |
| B09 + CYP12 | 31 | 69 | 54 | 24 | 44.5 | 20.8 |
| B09 + CYP13 | 61 | 51 | 32 | 42 | 46.5 | 12.4 |

**Table S6.** Extraction capacities of different fluorinate solvents.

| Solvent | Aristolochene | Valencene | Selinene | Vetispiradiene | Santalene | Bisabolol | Cadinol | Guaiene | Valerianol | Patchoulol |
|---|---|---|---|---|---|---|---|---|---|---|
| FC-40 | 15.24 | 17.44 | 15.64 | 17.54 | 17.44 | 31.94 | 28.54 | 29.94 | 27.74 | 30.54 |
| FC-770 | 14.24 | 16.44 | 14.64 | 16.54 | 16.44 | 30.94 | 27.54 | 28.94 | 26.74 | 29.54 |
| FC-3284 | 8.54 | 10.74 | 8.94 | 10.84 | 10.74 | 25.24 | 21.84 | 23.24 | 21.04 | 23.84 |
| CFL7160 | 14.04 | 16.24 | 14.44 | 16.34 | 16.24 | 30.74 | 27.34 | 28.74 | 26.54 | 29.34 |
| CXFL-3288 | 13.34 | 15.54 | 13.74 | 15.64 | 15.54 | 30.04 | 26.64 | 28.04 | 25.84 | 28.64 |
| CXFL-68 | 11.84 | 14.04 | 12.24 | 14.14 | 14.04 | 28.54 | 25.14 | 26.54 | 24.34 | 27.14 |
| CFL3000A | 10.74 | 12.94 | 11.14 | 13.04 | 12.94 | 27.44 | 24.04 | 25.44 | 23.24 | 26.04 |
| FC-43 | 13.04 | 15.24 | 13.44 | 15.34 | 15.24 | 29.74 | 26.34 | 27.74 | 25.54 | 28.34 |
| FC-72 | 9.44 | 11.64 | 9.84 | 11.74 | 11.64 | 26.14 | 22.74 | 24.14 | 21.94 | 24.74 |
| FC-3283 | 12.54 | 14.74 | 12.94 | 14.84 | 14.74 | 29.24 | 25.84 | 27.24 | 25.04 | 27.84 |

**Table S7.** GC-MS analysis of standard terpenoid mixture.

| Compound Name | Molecular formula | Classification | P (%) |
|---|---|---|---|
| Butanoic acid, 3-methyl-, 1-ethenyl-1,5-dimethyl-4-hexenyl ester | $C_{15}H_{26}O_2$ | Sesquiterpenoid | 73 |
| Camphene | $C_{10}H_{16}$ | Monoterpenoid | 88 |
| Bicyclo[3.1.1]heptane, 6,6-dimethyl-2-methylene-, (1S)- | $C_{10}H_{16}$ | Monoterpenoid | 78 |
| β-Myrcene | $C_{10}H_{16}$ | Monoterpenoid | 78 |
| Bicyclo[3.1.0]hex-2-ene, 4-methylene-1-(1-methylethyl)- | $C_{10}H_{14}$ | Monoterpenoid | 60 |
| Bicyclo[3.1.0]hex-2-ene, 2-methyl-5-(1-methylethyl)- | $C_{10}H_{16}$ | Monoterpenoid | 88 |
| 5-Isopropyl-2-methylbicyclo[3.1.0]hexan-2-ol # | $C_{10}H_{18}O$ | Monoterpenoid | 54 |
| p-Cymene | $C_{10}H_{14}$ | Monoterpenoid | 57 |
| Limonene | $C_{10}H_{16}$ | Monoterpenoid | 95 |
| Eucalyptol | $C_{10}H_{18}O$ | Monoterpenoid | 68 |
| 3-Carene | $C_{10}H_{16}$ | Monoterpenoid | 94 |
| γ-Terpinene | $C_{10}H_{16}$ | Monoterpenoid | 90 |
| Bicyclo[3.1.0]hexan-2-ol, 2-methyl-5-(1-methylethyl)-, (1α,2α,5α)- | $C_{10}H_{18}O$ | Monoterpenoid | 84 |
| Cyclohexene, 1-methyl-4-(1-methylethylidene)- | $C_{10}H_{18}O$ | Monoterpenoid | 92 |
| Fenchone | $C_{10}H_{18}O$ | Monoterpenoid | 95 |
| 3-Octanol, 3,7-dimethyl- | $C_{10}H_{18}O$ | Monoterpenoid | 95 |
| Fenchol | $C_{10}H_{18}O$ | Monoterpenoid | 88 |
| 1-Pentene, 5-(2,2-dimethylcyclopropyl)-2-methyl-4-methylene- | $C_{12}H_{20}$ | Ketone | 59 |
| 1,2-Dihydrolinalool | $C_{10}H_{18}O$ | Monoterpenoid | 84 |
| cis-Verbenol | $C_{10}H_{18}O$ | Monoterpenoid | 79 |
| (+)-2-Bornanone | $C_{10}H_{18}O$ | Monoterpenoid | 87 |
| Cyclohexanol, 5-methyl-2-(1-methylethenyl)- | $C_{10}H_{18}O$ | Monoterpenoid | 98 |
| Cyclohexanone, 5-methyl-2-(1-methylethyl)-, cis- | $C_{10}H_{18}O$ | Monoterpenoid | 94 |
| Isoborneol | $C_{10}H_{18}O$ | Monoterpenoid | 93 |

**Table S7.** Cont.

| Compound Name | Molecular formula | Classification | P (%) |
|---|---|---|---|
| Borneol | $C_{10}H_{18}O$ | Monoterpenoid | 95 |
| Cyclohexanol, 5-methyl-2-(1-methylethyl)-, [1S-(1α,2α,5β)]- | $C_{10}H_{20}O$ | Monoterpenoid | 76 |
| Terpinen-4-ol | $C_{10}H_{18}O$ | Monoterpenoid | 93 |
| α-Terpineol | $C_{10}H_{18}O$ | Monoterpenoid | 93 |
| Estragole | $C_{10}H_{12}O$ | Monoterpenoid | 80 |
| Bicyclo[3.1.1]hept-2-ene-2-methanol, 6,6-dimethyl- | $C_{10}H_{16}O$ | Monoterpenoid | 88 |
| Bicyclo[3.1.1]hept-3-en-2-one, 4,6,6-trimethyl- | $C_{10}H_{14}O$ | Monoterpenoid | 63 |
| 2,6-Octadiene, 1-(1-ethoxyethoxy)-3,7-dimethyl- | $C_{14}H_{26}O_2$ | Ketone | 89 |
| Cyclohexanone, 5-methyl-2-(1-methylethenyl)- | $C_{10}H_{16}O$ | Monoterpenoid | 85 |
| Carvone | $C_{10}H_{14}O$ | Monoterpenoid | 93 |
| Bicyclo[3.1.1]heptan-3-one, 2-hydroxy-2,6,6-trimethyl- | $C_{10}H_{16}O_2$ | Monoterpenoid | 53 |
| 2-Cyclohexen-1-one, 3-methyl-6-(1-methylethyl)- | $C_{10}H_{16}O$ | Monoterpenoid | 94 |
| Bicyclo[2.2.1]heptan-2-ol, 1,7,7-trimethyl-, acetate, (1S-endo)- | $C_{12}H_{20}O_2$ | Ketone | 89 |
| Phenol, 2-methyl-5-(1-methylethyl)- | $C_{10}H_{14}O$ | Monoterpenoid | 76 |
| Cyclohexanol, 5-methyl-2-(1-methylethyl)-, acetate, (1α,2α,5β)- | $C_{12}H_{22}O_2$ | Ketone | 84 |
| Phenol, 2-methyl-5-(1-methylethyl)- | $C_{10}H_{14}O$ | Monoterpenoid | 91 |
| Bicyclo[2.2.1]heptane-2,3-dione, 1,7,7-trimethyl-, (1S)- | $C_{10}H_{14}O_2$ | Monoterpenoid | 97 |
| Santolina alcohol | $C_{10}H_{18}O$ | Monoterpenoid | 93 |
| (1R,2R,3S,5R)-(-)-2,3-Pinanediol | $C10H_{18}O_2$ | Monoterpenoid | 91 |
| Bicyclo[2.2.1]heptane, 2-chloro-2,3,3-trimethyl- | $C_{10}H_{17}Cl$ | Monoterpenoid | 89 |
| 6-Octen-1-ol, 3,7-dimethyl-, acetate | $C_{12}H_{22}O_2$ | Ketone | 80 |
| Geranyl isovalerate | $C_{15}H_{26}O_2$ | Sesquiterpenoid | 81 |
| 4-Hexen-1-ol, 5-methyl-2-(1-methylethenyl)-, acetate | $C_{12}H_{20}O_2$ | Ketone | 100 |
| α-Damascone | $C_{13}H_{20}O$ | Ketone | 83 |

**Table S7.** Cont.

| Compound Name | Molecular formula | Classification | P (%) |
|---|---|---|---|
| 1,3-Cyclohexadiene, 5-(1,5-dimethyl-4-hexenyl)-2-methyl-, [S-(R*,S*)]- | $C_{15}H_{24}$ | Sesquiterpenoid | 64 |
| Caryophyllene | $C_{15}H_{24}$ | Sesquiterpenoid | 92 |
| cis-β-Farnesene | $C_{15}H_{24}$ | Sesquiterpenoid | 90 |
| Humulene | $C_{15}H_{24}$ | Sesquiterpenoid | 90 |
| 3-Buten-2-one, 4-(2,6,6-trimethyl-1-cyclohexen-1-yl)- | $C_{13}H_{20}O$ | Ketone | 85 |
| 1,3,6,10-Dodecatetraene, 3,7,11-trimethyl-, (Z,E)- | $C_{15}H_{24}$ | Sesquiterpenoid | 88 |
| 1,3,6,10-Dodecatetraene, 3,7,11-trimethyl-, (Z,E)- | $C_{15}H_{24}$ | Sesquiterpenoid | 85 |
| (E)-β-Farnesene | $C_{15}H_{24}$ | Sesquiterpenoid | 85 |
| Butylated Hydroxytoluene | $C_{15}H_{24}O$ | Sesquiterpenoid | 95 |
| 1-Isopropyl-4,7-dimethyl-1,2,3,5,6,8a-hexahydronaphthalene | $C_{15}5H_{24}$ | Sesquiterpenoid | 98 |
| 1,5-Diphenyl-2H-1,2,4-triazoline-3-thione | $C_{14}H_{11}N_3S$ | Ketone | 89 |
| (E)-β-Farnesene | $C_{15}H_{24}$ | Sesquiterpenoid | 95 |
| 1,6,10-Dodecatrien-3-ol, 3,7,11-trimethyl-, (E)- | $C_{15}H_{26}O$ | Sesquiterpenoid | 90 |
| Caryophyllene oxide | $C_{15}H_{24}O$ | Sesquiterpenoid | 95 |
| Cedrol | $C_{15}H_{26}O$ | Sesquiterpenoid | 84 |
| Cyclopentaneacetic acid, 3-oxo-2-(2-pentenyl)-, methyl ester, [1α,2α(Z)]- | $C_{13}H_{20}O_3$ | Ketone | 96 |
| α-Bisabolol | $C_{15}H_{26}O$ | Sesquiterpenoid | 97 |
| 2-Butenoic acid, 2-methyl-, 3,7-dimethyl-2,6-octadienyl ester, (E,Z)- | $C_{15}H_{24}O_2$ | Sesquiterpenoid | 79 |
| 1-Methylene-2b-hydroxymethyl-3,3-dimethyl-4b-(3-methylbut-2-enyl)-cyclohexane | $C_{15}H_{26}O$ | Sesquiterpenoid | 91 |
| Cyclopropane, 1-methyl-2-(3-methylpentyl)- | $C_{10}H_{20}$ | Monoterpenoid | 93 |
| Azulene, 1,4-dimethyl-7-(1-methylethyl)- | $C_{15}H_{18}$ | Sesquiterpenoid | 91 |
| Nootkatone | $C_{15}H_{22}O$ | Sesquiterpenoid | 81 |
| Neophytadiene | $C_{20}H_{38}$ | Diterpenoid | 95 |
| (E,E)-7,11,15-Trimethyl-3-methylene-hexadeca-1,6,10,14-tetraene | $C_{20}H_{32}$ | Diterpenoid | 88 |

**Table S7.** Cont.

| Compound Name | Molecular formula | Classification | P (%) |
|---|---|---|---|
| 3,7,11,15-Tetramethyl-2-hexadecen-1-ol | $C_{20}H_{40}O$ | Diterpenoid | 83 |
| 1-Methylene-2b-hydroxymethyl-3,3-dimethyl-4b-(3-methylbut-2-enyl)-cyclohexane | $C_{15}H_{26}O$ | Sesquiterpenoid | 60 |
| Musk ketone | $C_{15}H_{18}N_2O_5$ | Ketone | 95 |
| Phytol | $C_{20}H_{40}O$ | Diterpenoid | 93 |
| α-Santonin | $C_{15}H_{18}O_3$ | Sesquiterpenoid | 63 |
| 2-Methyl-3-(3-methyl-but-2-enyl)-2-(4-methyl-pent-3-enyl)-oxetane | $C_{15}H_{26}O$ | Sesquiterpenoid | 50 |
| Supraene | $C_{30}H_{50}$ | Triterpenoid | 88 |
| Squalene | $C_{30}H_{50}$ | Triterpenoid | 69 |

**Table S8.** GC-MS results of engineered *C. reinhardtii* strains.

| Compound Name | Molecular formula | Classification | P (%) |
|---|---|---|---|
| β-Vetispirene | $C_{15}H_{22}$ | Sesquiterpenoid | 65 |
| Cadinene | $C_{15}H_{22}O$ | Functionalized sesquiterpenoid | 69 |
| Nootkatone | $C_{15}H_{22}O$ | Functionalized sesquiterpenoid | 74 |
| Ylangenal | $C_{15}H_{22}O$ | Functionalized sesquiterpenoid | 38 |
| Valerianol | $C_{15}H_{22}O_2$ | Functionalized sesquiterpenoid | 74 |
| α-Bergamotene | $C_{15}H_{24}$ | Sesquiterpenoid | 60 |
| α-Cadinene | $C_{15}H_{24}$ | Sesquiterpenoid | 48 |
| α-Selinene | $C_{15}H_{24}$ | Sesquiterpenoid | 39 |
| Alloaromadendrene | $C_{15}H_{24}$ | Sesquiterpenoid | 39 |
| Aristolene | $C_{15}H_{24}$ | Sesquiterpenoid | 65 |
| Aristolochene | $C_{15}H_{24}$ | Sesquiterpenoid | 58 |
| Aromandendrene | $C_{15}H_{24}$ | Sesquiterpenoid | 52 |
| Copaene | $C_{15}H_{24}$ | Sesquiterpenoid | 71 |
| Germacrene D | $C_{15}H_{24}$ | Sesquiterpenoid | 80 |
| γ-Gurjunene | $C_{15}H_{24}$ | Sesquiterpenoid | 48 |
| Humulene | $C_{15}H_{24}$ | Sesquiterpenoid | 37 |
| Muurolane | $C_{15}H_{24}$ | Sesquiterpenoid | 63 |
| Rotundene | $C_{15}H_{24}$ | Sesquiterpenoid | 78 |
| Valencene | $C_{15}H_{24}$ | Sesquiterpenoid | 87 |
| α-Bergamotene | $C_{15}H_{24}$ | Sesquiterpenoid | 42 |
| α-Guaiene | $C_{15}H_{24}$ | Sesquiterpenoid | 68 |
| β-Cadinene | $C_{15}H_{24}$ | Sesquiterpenoid | 52 |
| β-Guaiene | $C_{15}H_{24}$ | Sesquiterpenoid | 91 |
| β-Selinene | $C_{15}H_{24}$ | Sesquiterpenoid | 57 |
| γ-Muurolene | $C_{15}H_{24}$ | Sesquiterpenoid | 62 |
| δ-Guaiene | $C_{15}H_{24}$ | Sesquiterpenoid | 87 |
| α-Muurolene | $C_{15}H_{24}O$ | Functionalized sesquiterpenoid | 42 |

**Table S8.** Cont.

| Compound Name | Molecular formula | Classification | P (%) |
|---|---|---|---|
| α-Santalol | $C_{15}H_{24}O$ | Functionalized sesquiterpenoid | 59 |
| Alloaromadendrene oxide | $C_{15}H_{24}O$ | Functionalized sesquiterpenoid | 81 |
| Aromadendrene oxide-(1) | $C_{15}H_{24}O$ | Functionalized sesquiterpenoid | 69 |
| Humulene epoxide I | $C_{15}H_{24}O$ | Functionalized sesquiterpenoid | 54 |
| Humulene oxide II | $C_{15}H_{24}O$ | Functionalized sesquiterpenoid | 66 |
| α-Agarofuran | $C_{15}H_{24}O$ | Functionalized sesquiterpenoid | 38 |
| α-Bisabolene epoxide | $C_{15}H_{24}O$ | Functionalized sesquiterpenoid | 45 |
| β-Santalol | $C_{15}H_{24}O$ | Functionalized sesquiterpenoid | 74 |
| α-Eudesmol | $C_{15}H_{26}O$ | Functionalized sesquiterpenoid | 60 |
| Agarospirol | $C_{15}H_{26}O$ | Functionalized sesquiterpenoid | 48 |
| Cubedol | $C_{15}H_{26}O$ | Functionalized sesquiterpenoid | 39 |
| Epiglobulol | $C_{15}H_{26}O$ | Functionalized sesquiterpenoid | 28 |
| Globulol | $C_{15}H_{26}O$ | Functionalized sesquiterpenoid | 65 |
| Guaiol | $C_{15}H_{26}O$ | Functionalized sesquiterpenoid | 58 |
| Patchouli alcohol | $C_{15}H_{26}O$ | Functionalized sesquiterpenoid | 51 |
| α-Bisabolol | $C_{15}H_{26}O$ | Functionalized sesquiterpenoid | 44 |

**Table S9.** Mass spectra of sesquiterpenoids identified by GC – MS.

| Compound Name | Molecular formula | Structure | Mass spectra |
|---|---|---|---|
| Santalene | $C_{15}H_{24}$ | 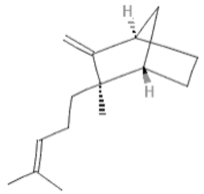 | 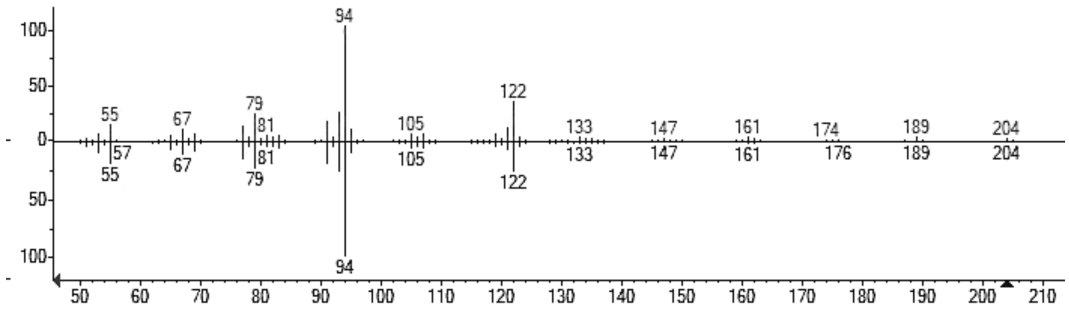 |
| Bergamotene | $C_{15}H_{24}$ | 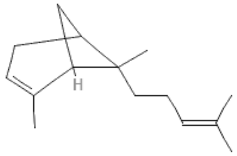 | 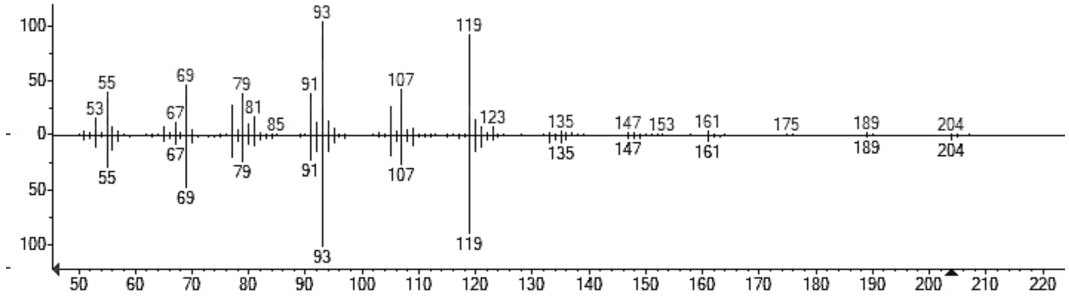 |
| α-Santalol | $C_{15}H_{24}O$ | 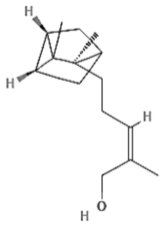 | 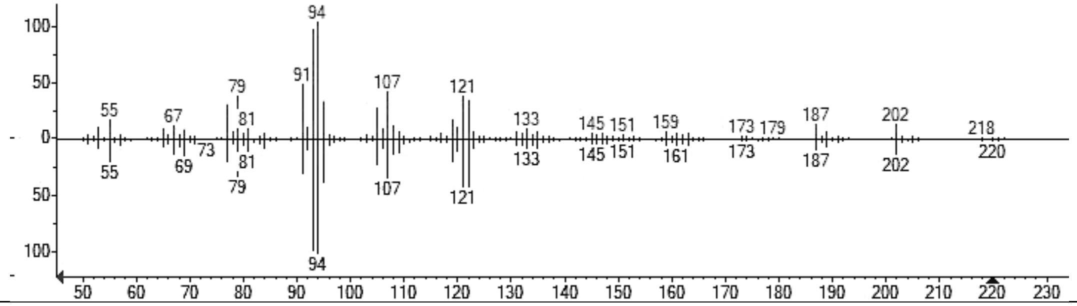 |

**Table S9.** Cont.

| Compound Name | Molecular formula | Structure | Mass spectra |
|---|---|---|---|
| Cedrene | $C_{15}H_{24}$ | 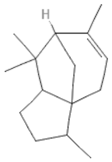 | 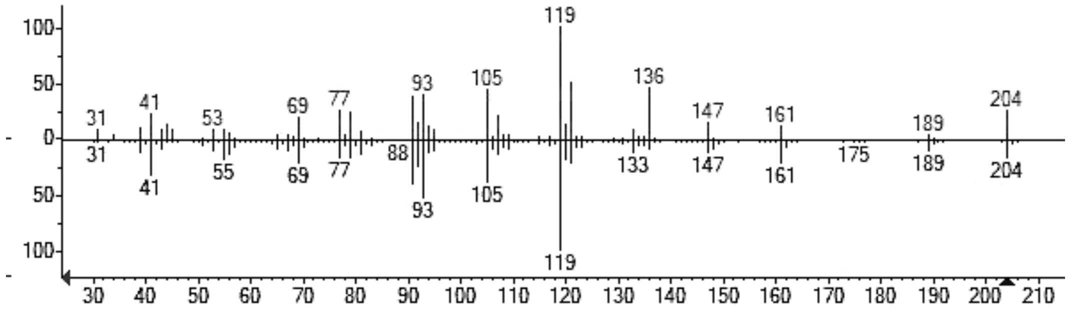 |
| Curcumene | $C_{15}H_{22}$ | 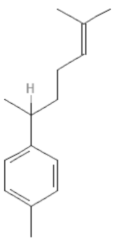 | 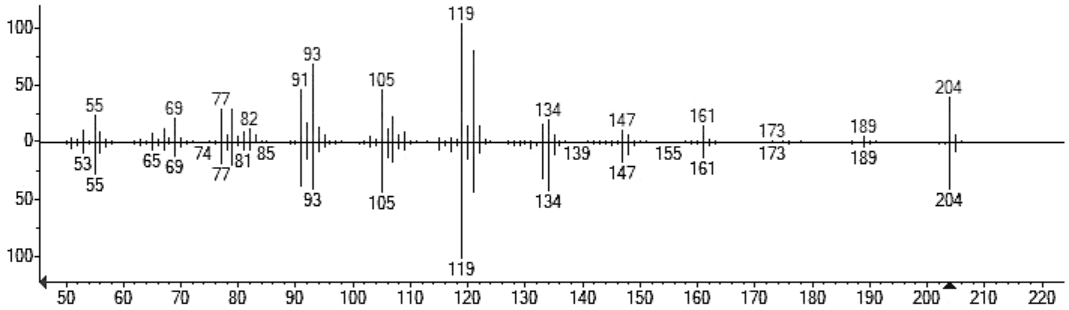 |
| Himalachene | $C_{15}H_{24}$ | 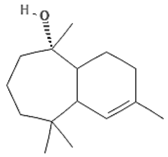 | 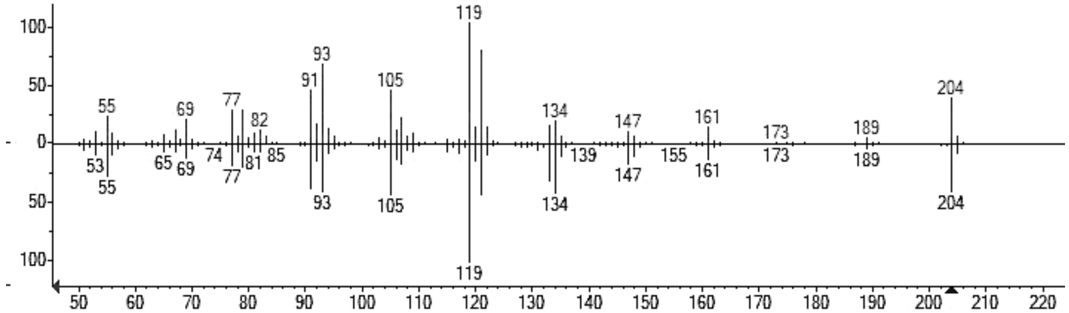 |

**Table S9.** Cont.

| Compound Name | Molecular formula | Structure | Mass spectra |
|---|---|---|---|
| Isoledene | $C_{15}H_{24}$ | 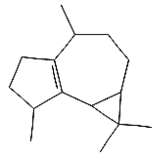 | 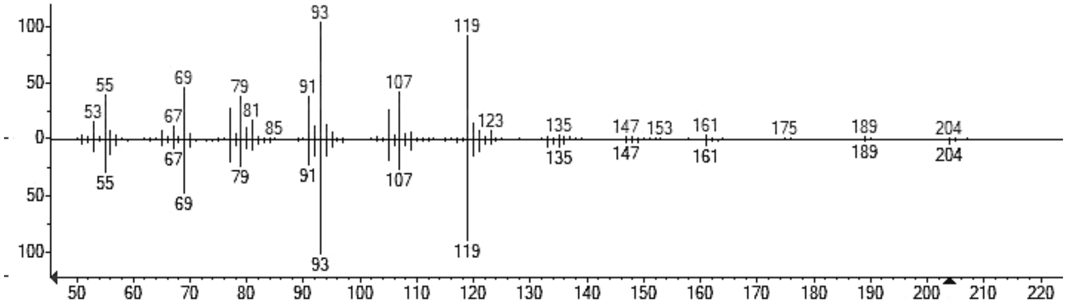 |
| Longipinene | $C_{15}H_{24}$ | 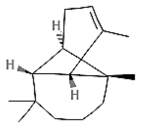 | 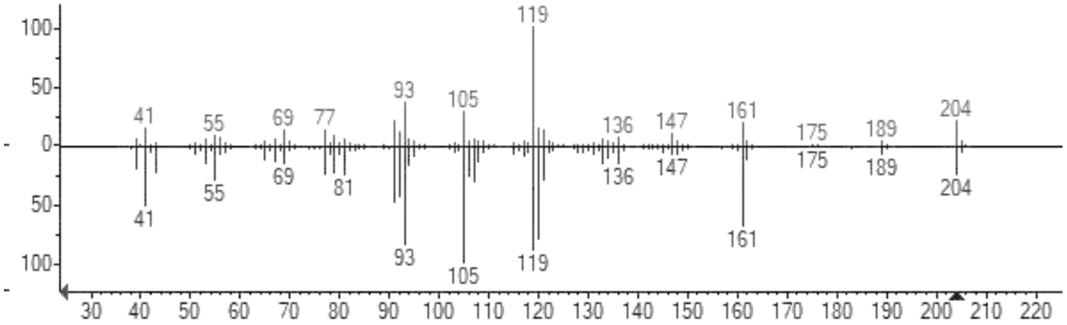 |
| Germacrene D | $C_{15}H_{24}$ | 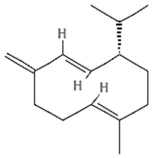 | 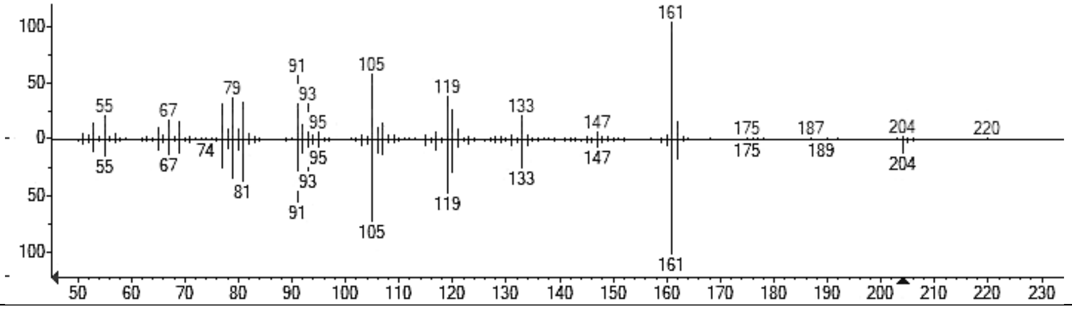 |

**Table S9.** Cont.

| Compound Name | Molecular formula | Structure | Mass spectra |
|---|---|---|---|
| γ-Muurolene | $C_{15}H_{24}$ | 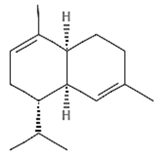 | 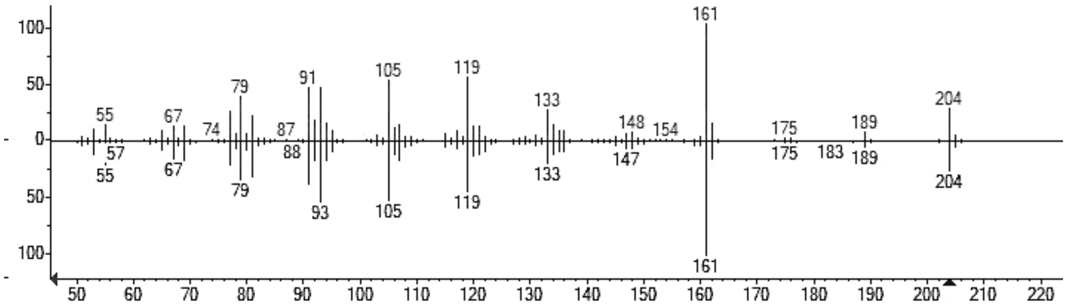  |
| β-Agarofuran | $C_{15}H_{26}O$ | 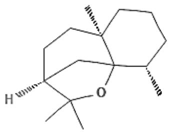 | 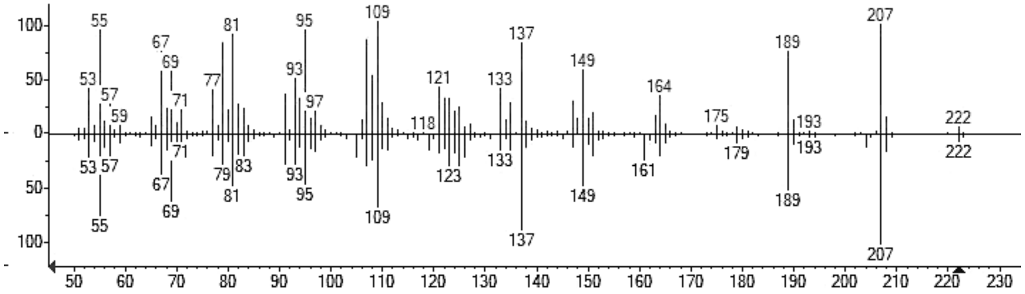  |
| α-Humulene | $C_{15}H_{24}$ | 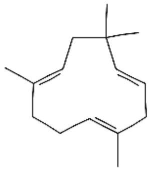 | 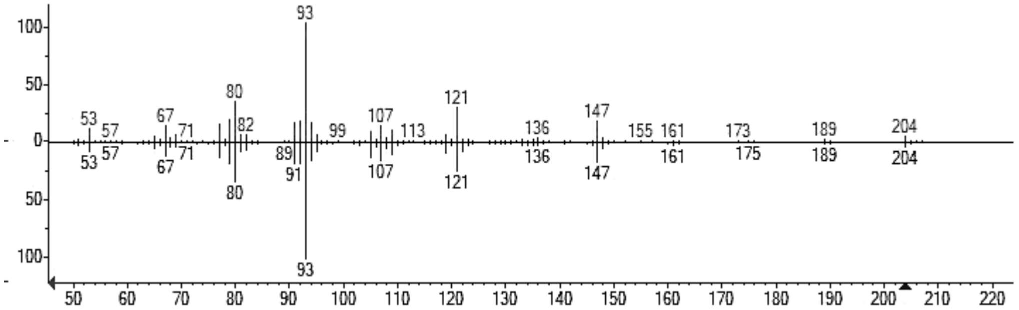  |

**Table S9.** Cont.

| Compound Name | Molecular formula | Structure | Mass spectra |
|---|---|---|---|
| α-Guaiene | $C_{15}H_{24}$ | 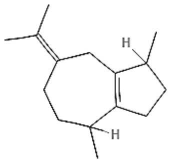 | 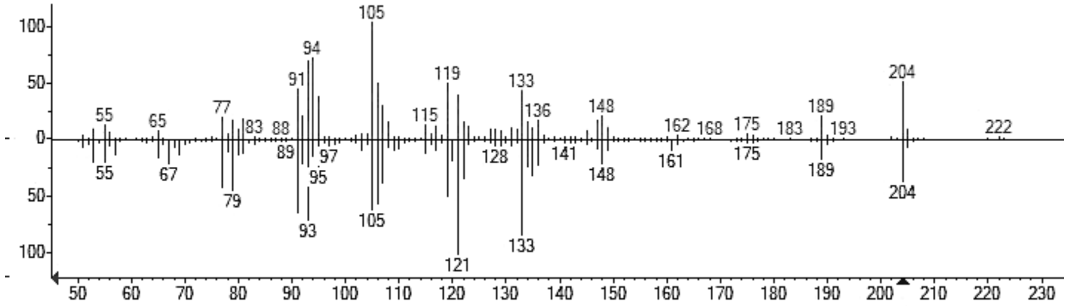 |
| Copaene | $C_{15}H_{24}$ | 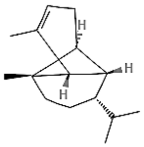 | 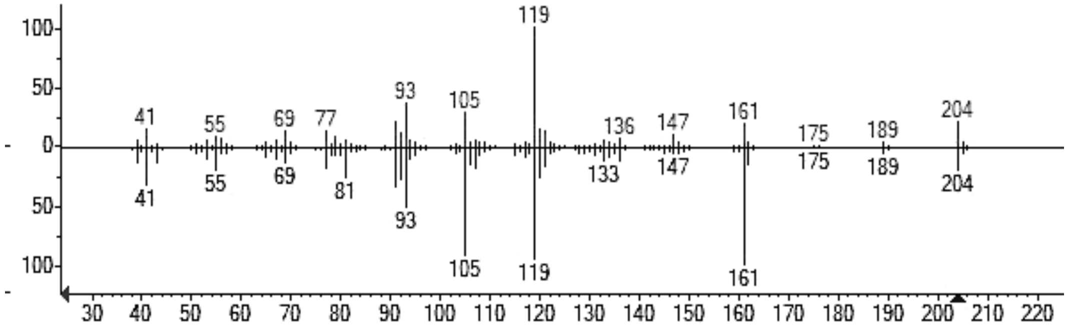 |
| Aromadendrene | $C_{15}H_{24}$ | 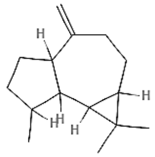 | 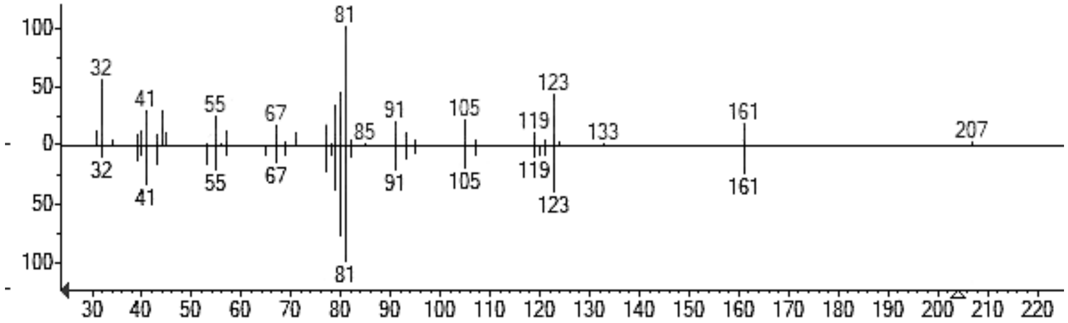 |

**Table S9.** Cont.

| Compound Name | Molecular formula | Structure | Mass spectra |
|---|---|---|---|
| β-Bourbonene | $C_{15}H_{24}$ | 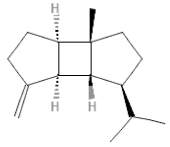 | 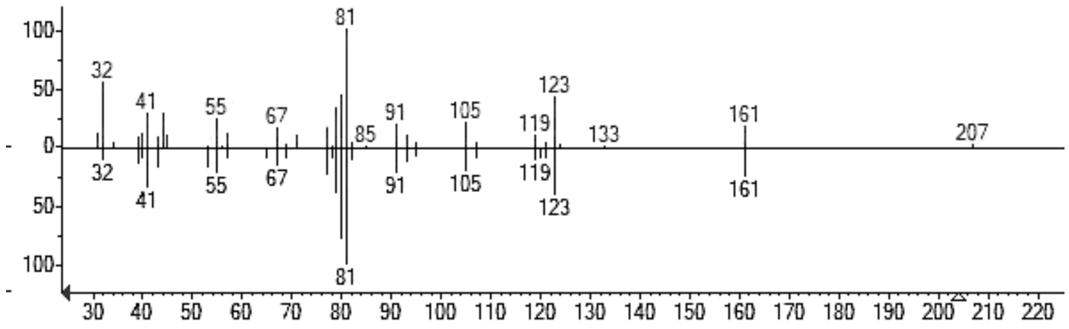 |
| α-Bourbonene | $C_{15}H_{24}$ | 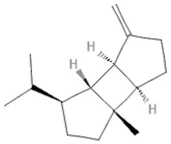 | 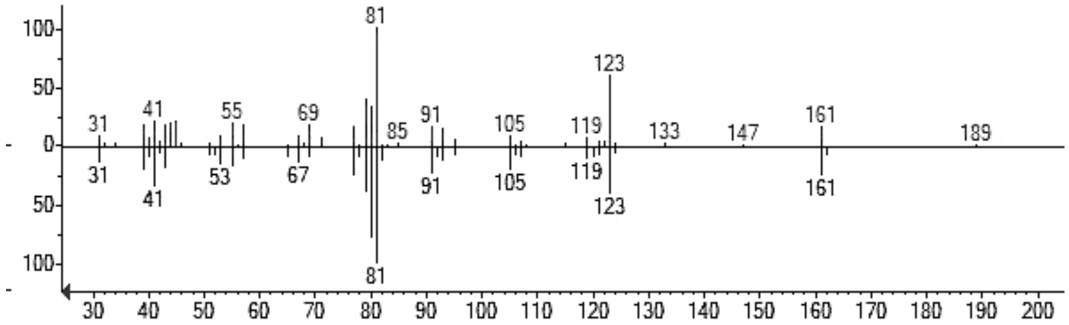 |
| Alloaromadendrene | $C_{15}H_{24}$ | 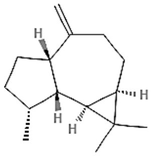 | 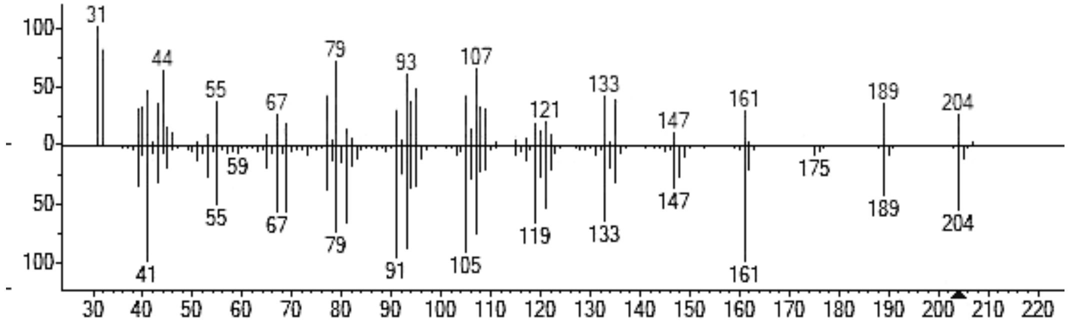 |

**Table S9.** Cont.

| Compound Name | Molecular formula | Structure | Mass spectra |
|---|---|---|---|
| Seychellene | $C_{15}H_{24}$ | 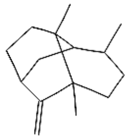 | 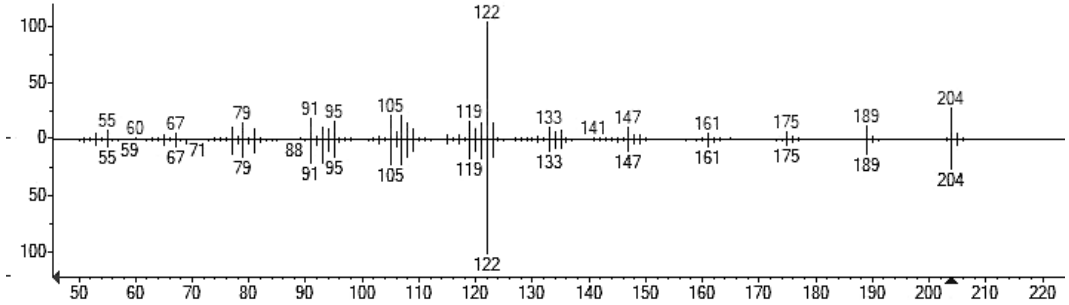 |
| Patchoulene | $C_{15}H_{24}$ | 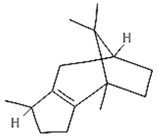 | 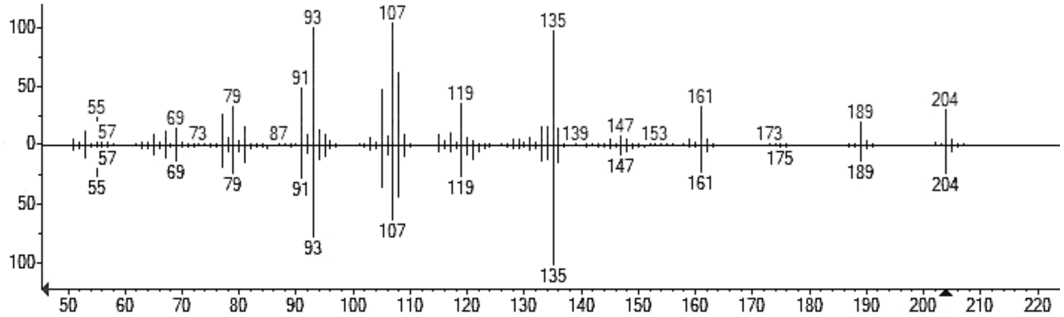 |
| Cadinol | $C_{15}H_{26}O$ | 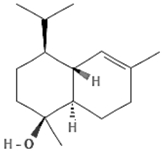 | 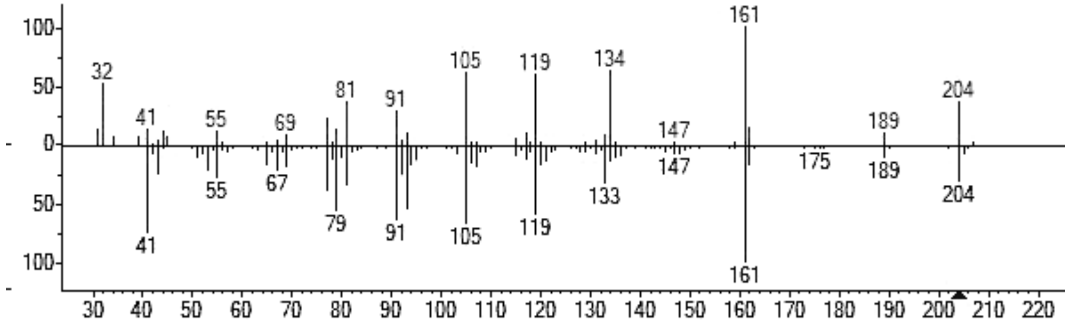 |

**Table S9.** Cont.

| Compound Name | Molecular formula | Structure | Mass spectra |
|---|---|---|---|
| Selinene | $C_{15}H_{24}$ | 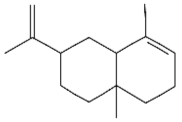 | 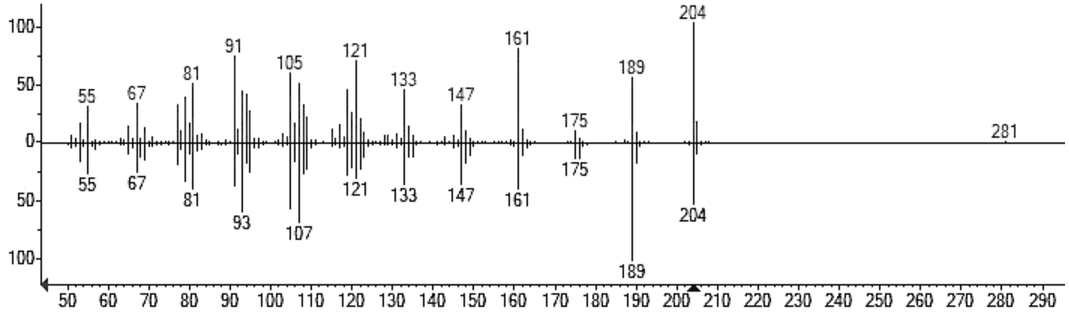 |
| Eremophilane | $C_{15}H_{24}$ | 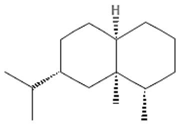 | 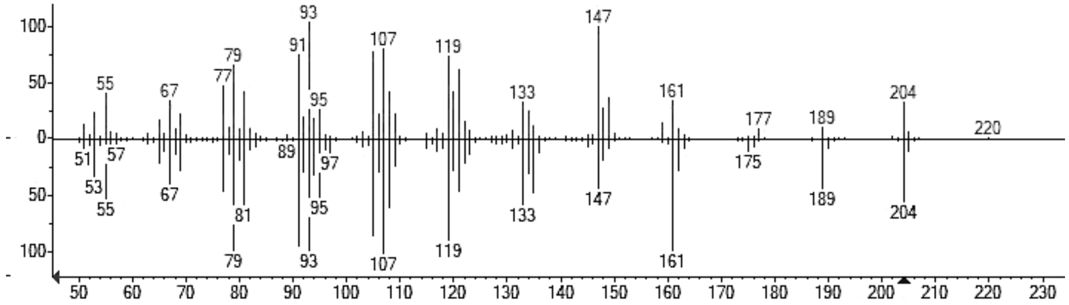 |
| δ-Guaiene | $C_{15}H_{24}$ | 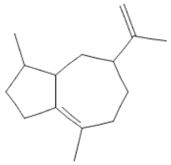 | 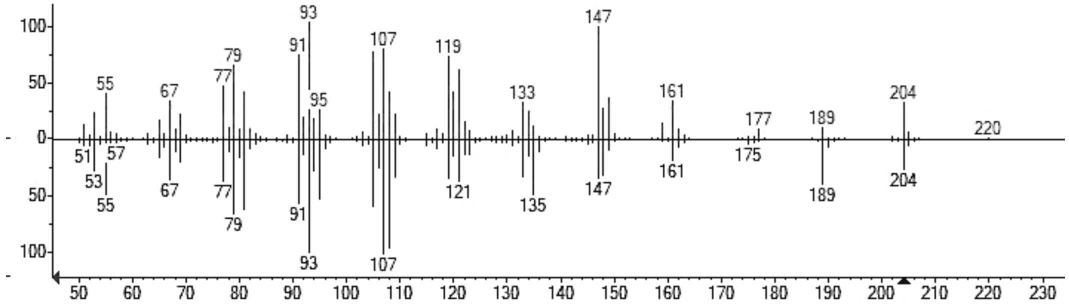 |

**Table S9.** Cont.

| Compound Name | Molecular formula | Structure | Mass spectra |
|---|---|---|---|
| Aristolochene | $C_{15}H_{24}$ | 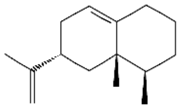 | 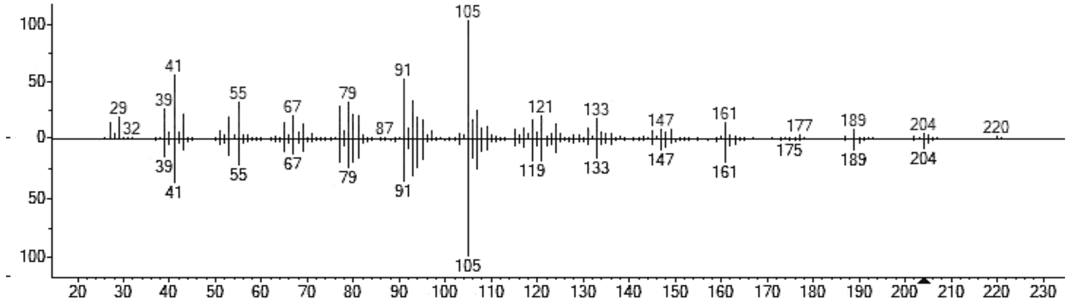 |
| Valerianol | $C_{15}H_{26}O$ | 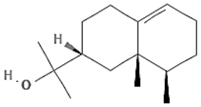 | 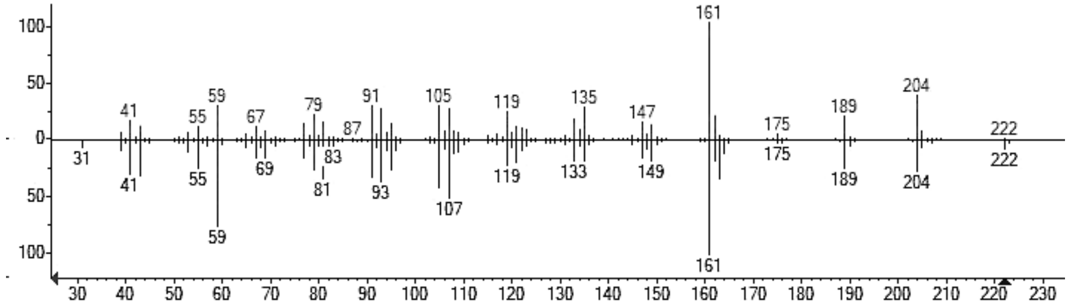 |
| Patchoulol | $C_{15}H_{26}O$ | 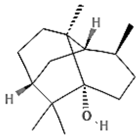 | 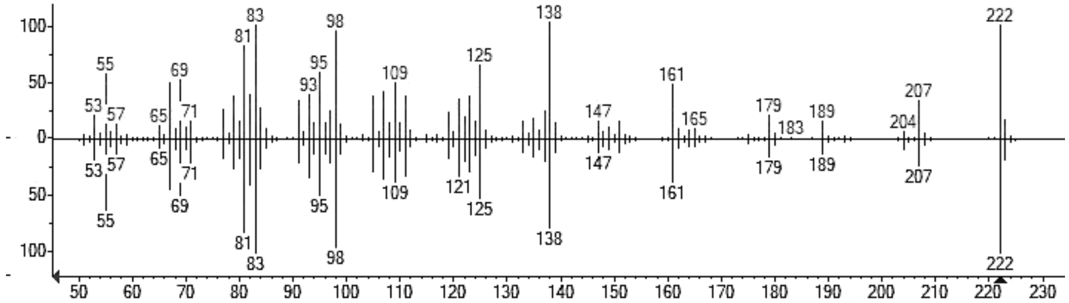 |

**Table S9.** Cont.

| Compound Name | Molecular formula | Structure | Mass spectra |
|---|---|---|---|
| Bisabolol | $C_{15}H_{26}O$ | 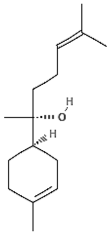 | 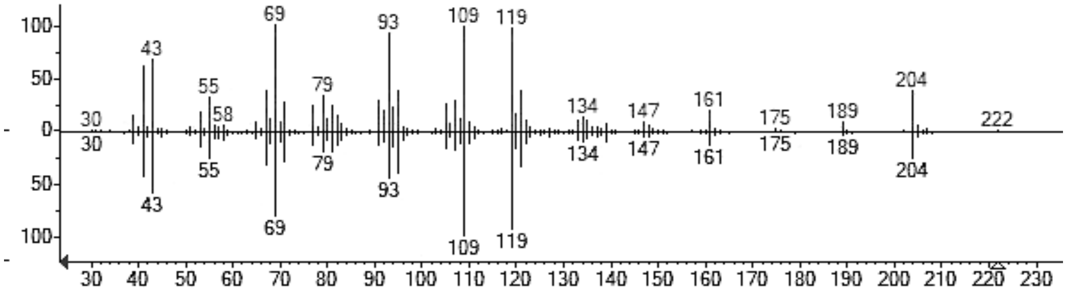  |
| Valencene | $C_{15}H_{24}$ | 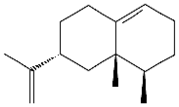 | 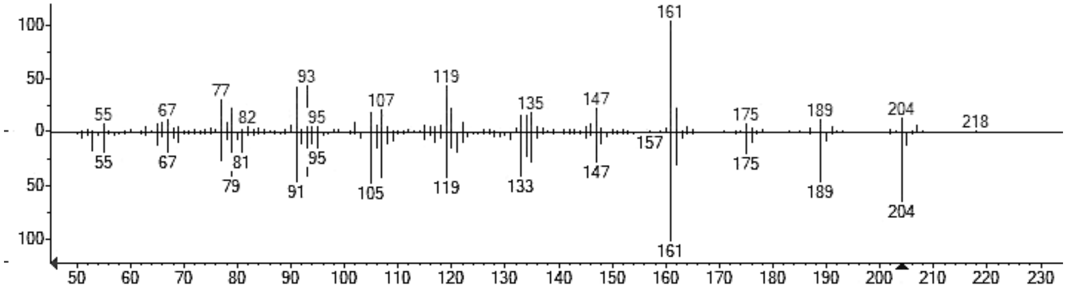  |
| Gurjunene | $C_{15}H_{24}$ | 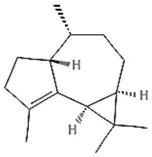 | 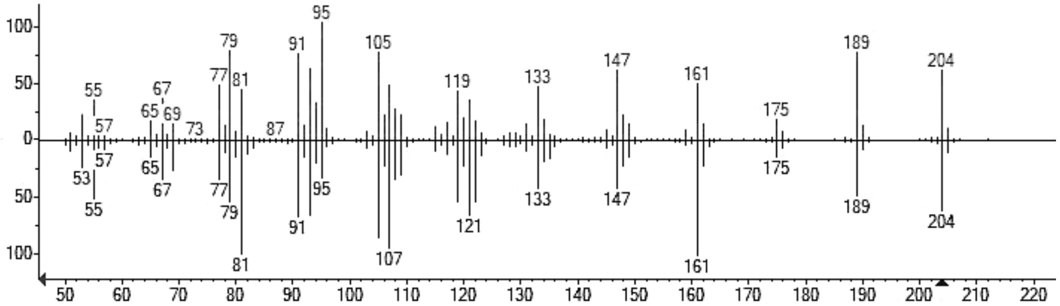  |

**Table S9.** Cont.

| Compound Name | Molecular formula | Structure | Mass spectra |
|---|---|---|---|
| α-Santalol | $C_{15}H_{24}O$ | 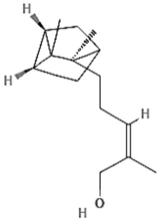 | 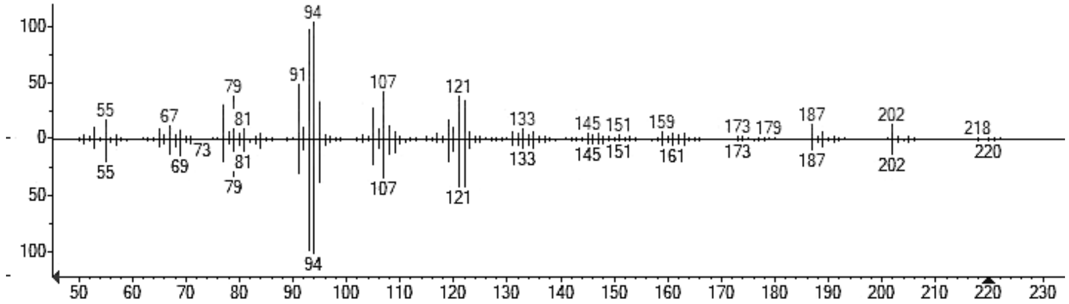 |
| β-Santalol | $C_{15}H_{24}O$ | 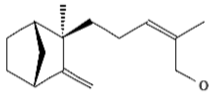 | 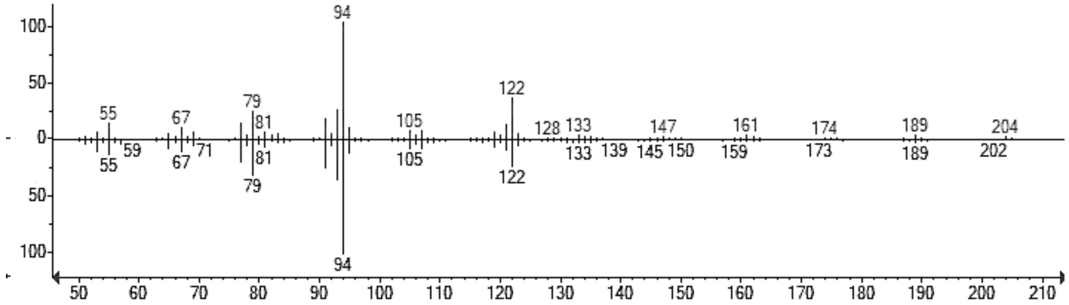 |
| Isolongifolol | $C_{16}H_{28}O$ | 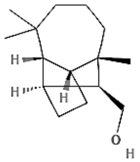 | 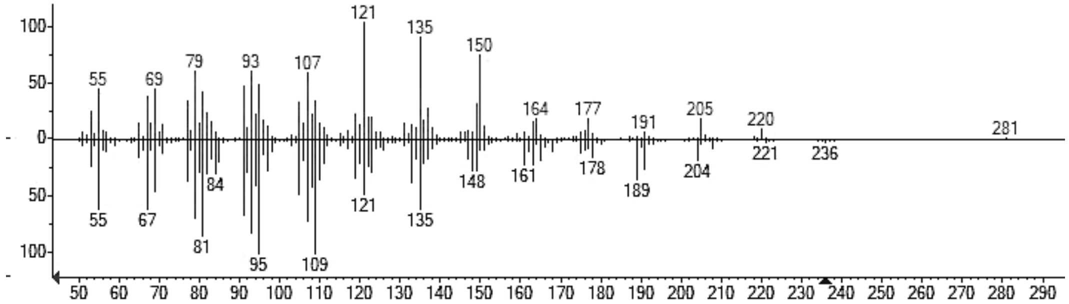 |

**Table S9.** Cont.

| Compound Name | Molecular formula | Structure | Mass spectra |
|---|---|---|---|
| Cubenol | $C_{15}H_{26}O$ | 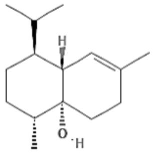 | 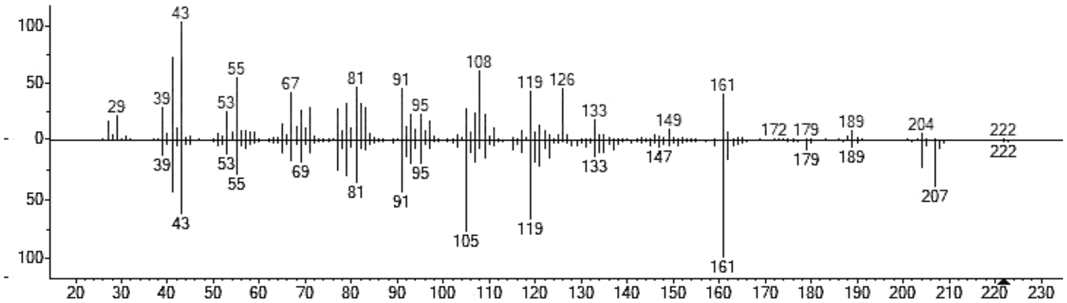 |
| Khusimol | $C_{15}H_{24}O$ | 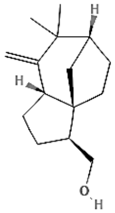 | 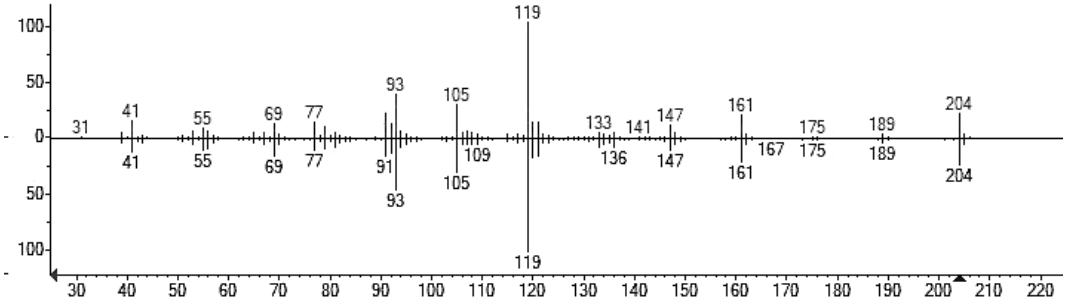 |
| Cadinene | $C_{15}H_{24}$ | 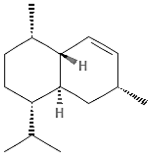 | 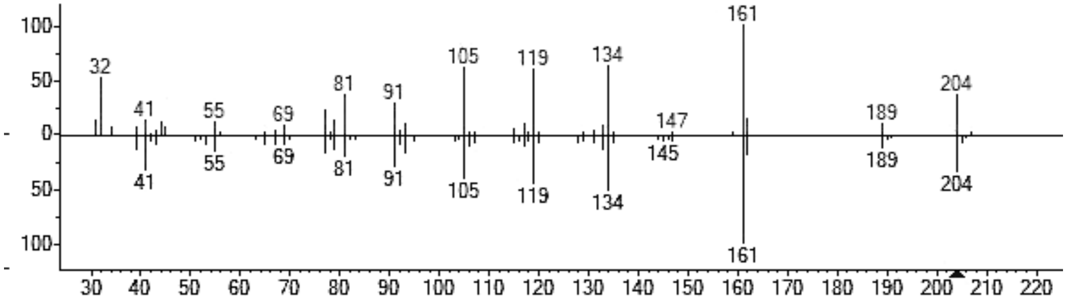 |

**Table S9.** Cont.

| Compound Name | Molecular formula | Structure | Mass spectra |
|---|---|---|---|
| Guaiol | $C_{15}H_{26}O$ | 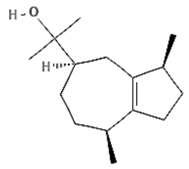 | 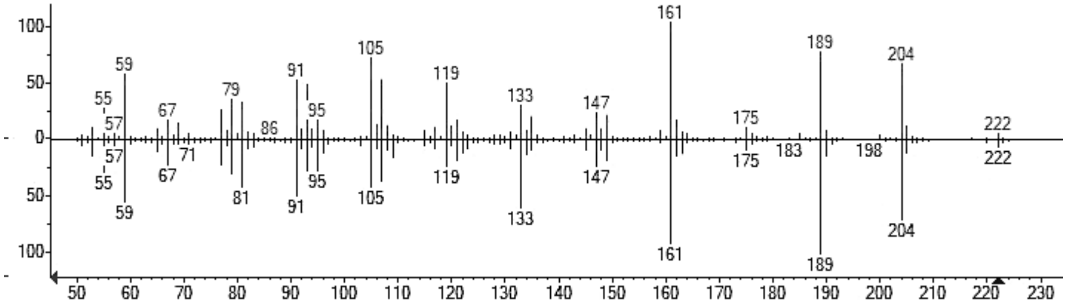 |
| Globulol | $C_{15}H_{26}O$ | 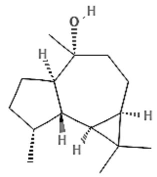 | 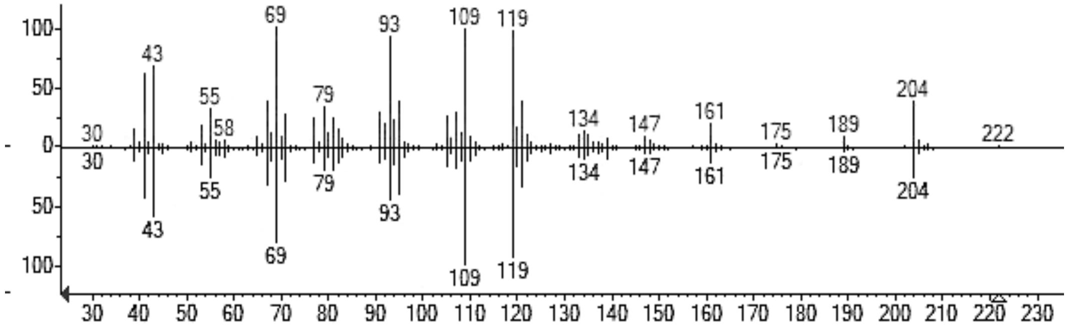 |
| Guaia-1(10),11-dien-15,2-olide | $C_{15}H_{20}O_2$ | 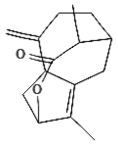 | 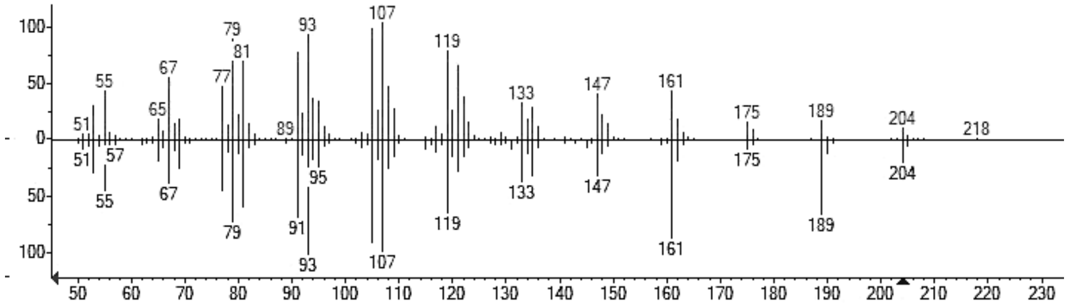 |

**Table S9.** Cont.

| Compound Name | Molecular formula | Structure | Mass spectra |
|---|---|---|---|
| Eudesmol | $C_{15}H_{26}0$ | 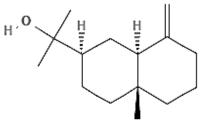 | 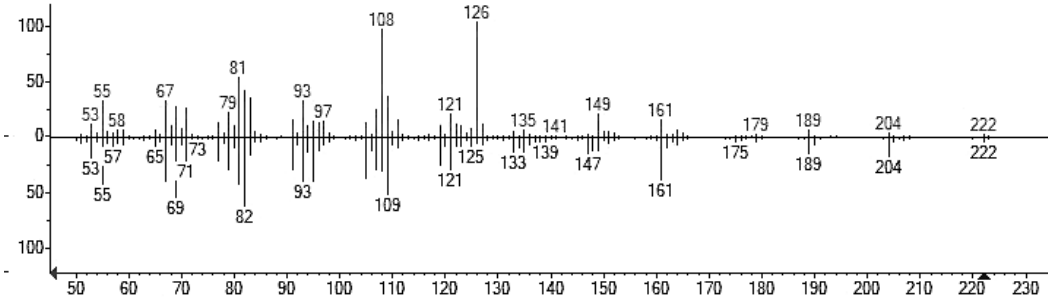 |
| Humulane-1,6-dien-3-ol | $C_{15}H_{26}O$ | 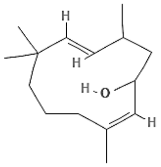 | 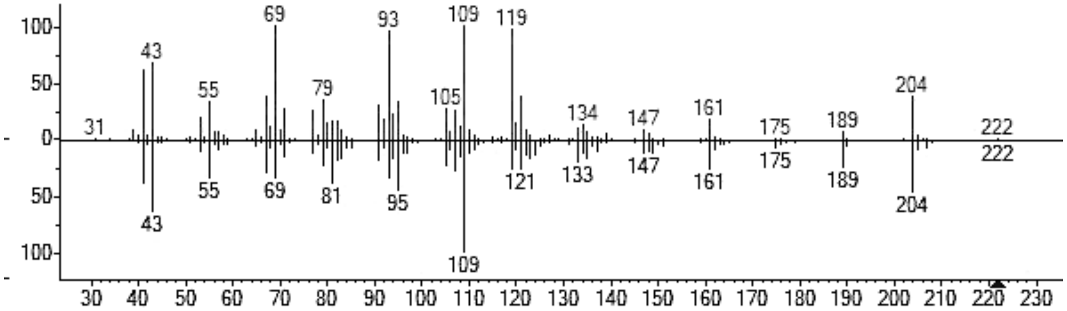 |
| Bisabolol oxide B | $C_{15}H_{26}O_2$ | 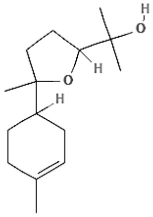 | 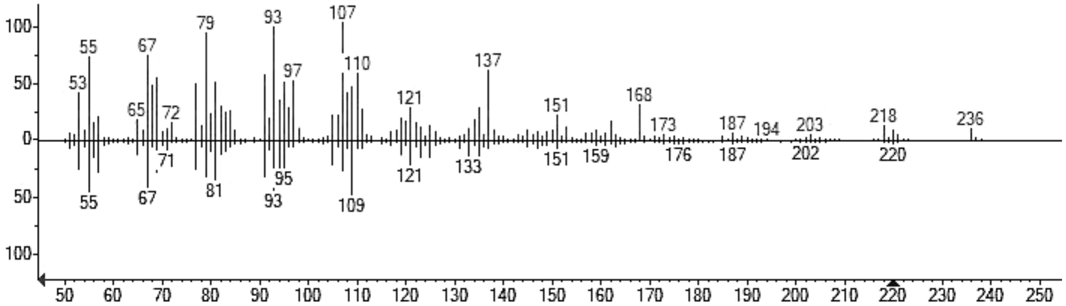 |

**Table S9.** Cont.

| ompound Name | Molecular formula | Structure | Mass spectra |
|---|---|---|---|
| Bisabolol oxide A | $C_{15}H_{26}O_2$ | 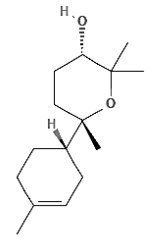 | 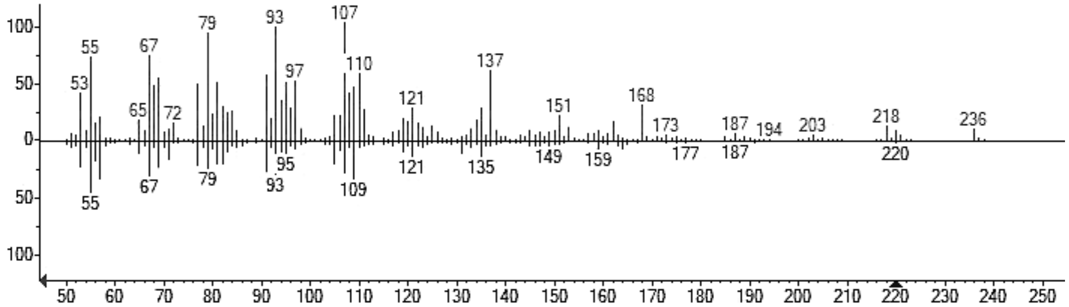  |
| Ylangene | $C_{15}H_{24}$ | 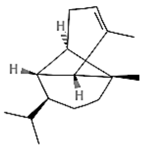 | 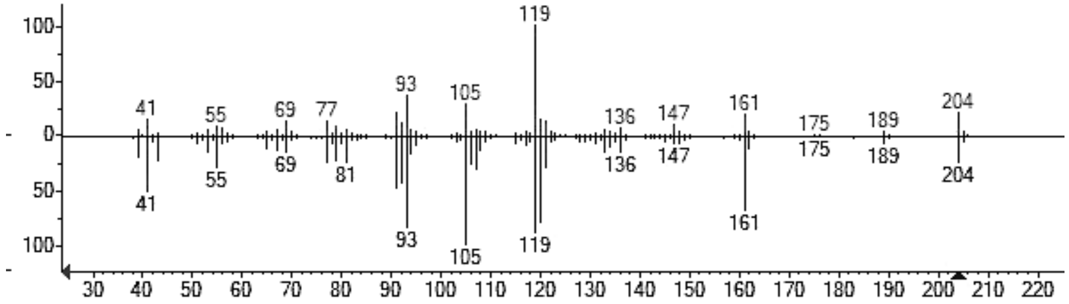  |
| Agidol | $C_{15}H_{24}O$ | 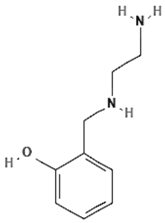 | 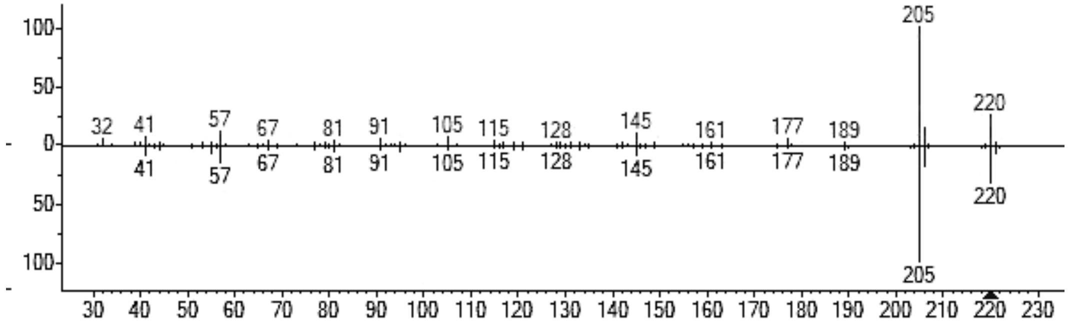  |

**Table S9.** Cont.

| Compound Name | Molecular formula | Structure | Mass spectra |
|---|---|---|---|
| Ledol | $C_{15}H_{26}O$ | 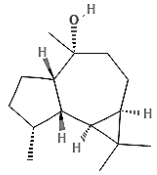 | 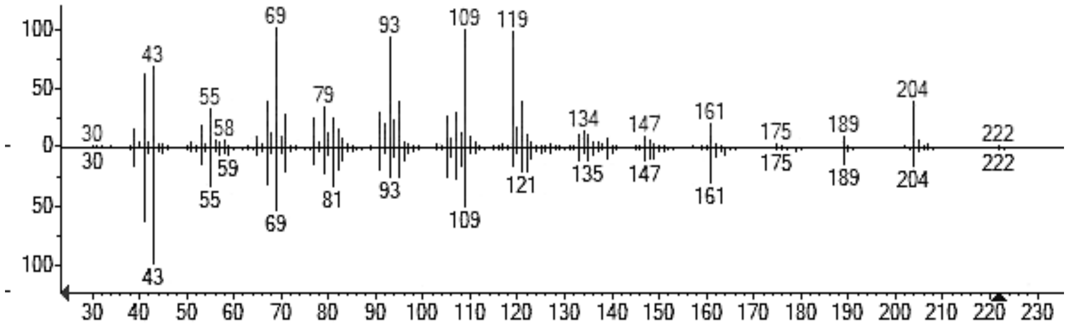 |
| Aromadendrene oxide | $C_{15}H_{24}O$ | 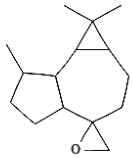 | 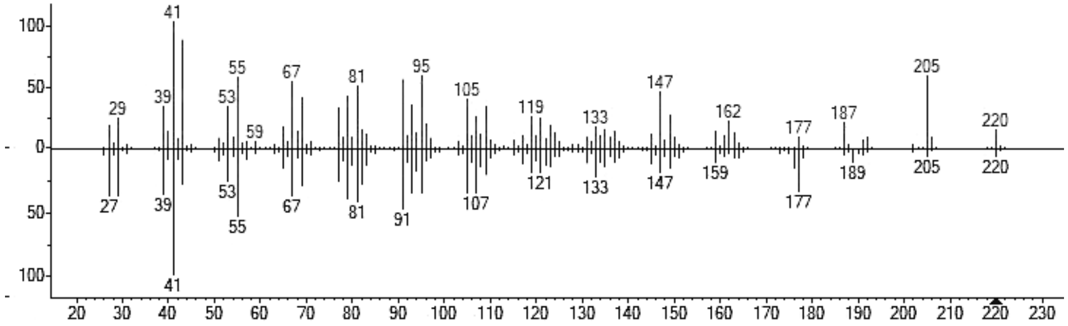 |

#### SI Files

Files can be accessed via a temporary link (https://rb.gy/lpdr3a).

**File S1.** Genetic constructs used in this study.
(Sequences and maps of plasmids used for *C. reinhardtii* transformation)

**File S2.** GC-MS/FID chromatograms for *C. reinhardtii* strains harboring plasmids A.
(Raw chromatographic data for strains expressing plasmids A)

**File S3.** GC-MS/FID chromatograms for *C. reinhardtii* strains harboring plasmids B.
(Raw chromatographic data for strains expressing plasmids B)

**File S4.** GC-MS/FID chromatograms for *C. reinhardtii* strains harboring plasmids C.
(Raw chromatographic data for strains expressing plasmids C)

**File S5.** GC-MS/FID chromatograms for *C. reinhardtii* strains co-expressing plasmids B and CYP.
Raw chromatographic data for strains expressing both plasmids B and cytochrome P450 enzymes.

**File S6.** GC-MS/FID chromatograms for *C. reinhardtii* strains co-expressing plasmids B and CYP grown with different carbon sources.
(Comparison of sesquiterpenoid production under various growth conditions)

**File S7.** Raw GC-MS/FID data.
(Raw data from GC-MS/FID instrument)
